# Web engine for tumor pathology image retrievals on massive scales

**DOI:** 10.1101/2025.10.25.684566

**Authors:** Zisha Zhong, Jinlin Huang, Xinjing Li, Lijuan Song, Lanqi Gong, Beibei Ru, Rohit Paul, Jason Levine, Yu Zhang, Kenneth Aldape, Peng Jiang

**Affiliations:** Cancer Data Science Lab, Center for Cancer Research, National Cancer Institute, National Institutes of Health; Bethesda, Maryland 20892, U.S.A; Department of Pathology, Sun Yat-sen University Cancer Center; State Key Laboratory of Oncology in South China, Guangzhou, Guangdong, P.R. China; Department of Pathology, Longyan First Affiliated Hospital of Fujian Medical University, Longyan, Fujian, P.R. China; Department of Pathology, Sun Yat-sen University Cancer Center Gansu Hospital, Lanzhou, Gansu, P.R. China; Office of the Director, Center for Cancer Research, National Cancer Institute, National Institutes of Health, Bethesda, MD, U.S.A; Sun Yat-sen University Cancer Center; State Key Laboratory of Oncology in South China; Collaborative Innovation Center for Cancer Medicine, Guangzhou, Guangdong, P.R. China; Laboratory of Pathology, Center for Cancer Research, National Cancer Institute, National Institutes of Health; Bethesda, Maryland 20892, U.S.A

## Abstract

Hematoxylin and Eosin staining (H&E) is widely used in clinical practice, but efficient and versatile image retrieval tools are lacking. We developed the H&E Retrieval Engine (HERE, https://hereapp.ccr.cancer.gov) to analyze patient cases based on image similarities to database records. Using H&E image regions as input, HERE searches 21.2 terabytes of whole-slide images from multiple tumor histopathology cohorts through a 12.1-gigabyte memory index, and returns top images containing regions similar to the query. HERE scans high-resolution images in the database using accurate artificial intelligence encoding and ultra-efficient hierarchical skip indexing. HERE demonstrated performance superior to existing image retrieval tools based on blinded pathologist scoring using benchmark queries that represent key image features of human tumors. By pairing spatial transcriptomics with H&E images, HERE also enables retrieving image features from gene transcriptomics input and identifies molecular pathways associated with tumor histologies.

## Introduction

For over 100 years, Hematoxylin and Eosin (H&E) staining has been the most widely used tissue staining approach for clinical diagnosis and histopathology, partly due to its ease, speed, and low cost. As a result, H&E images are abundant, and image databases contain the vast morphological complexity derived from a variety of clinical applications. The H&E image datasets from the Cancer Genome Atlas (TCGA) project and the Lab of Pathology at the National Cancer Institute (NCI) now each exceed seven terabytes. Despite many previous efforts to manage and retrieve H&E images based on visual features using content-based image retrieval (CBIR)^1–24^, we are not aware of any public web portal for clinicians and researchers to search image databases on a terabyte-scale and above, analogous to BLAST^25^ for sequencing data.

For a given H&E image database, retrieval engines would ideally (1) index the entire image collection and (2) search the entirety of each image, covering all local regions without down-sampling. We are aware of two open-access web portals that provide image search capabilities on a limited scale: Luigi^26^ and PLIP^27^. A single whole-slide image usually contains millions or even billions of pixels, resulting in a file size of several gigabytes and presenting a challenge for search functions. To address this challenge, Luigi selects three or more representative patches for each slide to reduce the search space^26^. Similarly, other CBIR methods with available source code (but not web portals), such as SISH^21^, Yottixel^19^, and RetCCL^23^, select a small fraction of representative patches from each whole-slide image to create a “mosaic” retrieval space. Rather than selecting patches, PLIP resizes the query image into a 224 x 224-pixel image to compute one overall similarity between the input and query images^27^. However, a whole slide image may cover heterogeneous tissue regions. None of these frameworks preserve image continuity and resolution.

One reason for the lack of large-scale H&E retrieval engines is algorithm scalability. Current CBIR frameworks generally operate in three steps. First, the framework encodes images or local patches as feature vectors through neural network encoders, such as deep convolutional networks or transformers. Second, most frameworks hash feature vectors into binary codes to enable rapid comparisons^3,7–9,12,24^. Third, many frameworks further organize feature vectors into structured indices to accelerate searching^14,17,19,21^. However, these solutions may fail for large-scale image data. For example, the SMILY framework used a k-dimensional (KD) tree to organize image vectors^14^. However, modern CBIR frameworks encode image features into ultra-high dimensions, and the speed performance advantages afforded by KD trees deteriorate in high-dimensional feature space due to the curse-of-dimensionality problem^28^. To address this, the SISH framework^21^ organized feature vectors through the Van Emde Boas (vEB) tree^29^. However, the vEB tree has a prohibitive memory space requirement, making it unsuitable for indexing a large amount of images on typical computer servers.

Another limitation of current H&E search tools is the lack of retrieval functions that bridge genetic and molecular information with image features. Molecular biology approaches have driven many cancer research and therapeutics breakthroughs^30^. Clinical pathology diagnoses depend heavily on molecular markers. However, H&E images contain only morphological and structural information, and we are unaware of an existing tool that enables image retrievals using genetic or molecular information as query input.

We present the H&E Retrieval Engine (HERE, https://hereapp.ccr.cancer.gov) that retrieves whole-slide images from the entire collection at the NCI Lab of Pathology, TCGA, and Clinical Proteomic Tumor Analysis Consortium (CPTAC) within seconds, along with a framework for others to develop a retrieval engine on any H&E data. We utilized optimized artificial intelligence (AI) encoding and hierarchical skip indexing of image features to enable rapid searching across terabytes of images with a desktop computer. To evaluate the retrieval accuracy of web portals, we established the HERE101 benchmark, covering diverse image features found in human cancers. Evaluations by pathologists demonstrated the reliable and superior performance of HERE relative to existing retrieval tools. In addition to image-to-image searches, HERE allows gene transcriptomics-to-image search functions derived from spatial transcriptomics data paired with H&E images. This functionality can identify the top molecular pathways associated with tumor histologies, such as complement activation. Using in-vivo mouse models, we showed that overexpressing an inhibitor of complement activation suppressed tumor growth.

## Results

### Overview of the HERE framework

The HERE framework includes two modules: engine development (Fig. 1a) and online retrieval (Fig. 1b). The development of an H&E retrieval engine depends on the accurate encoding of image patches and regions into feature vectors (Fig. 1a and Extended Data Fig. 1 - Stage 1) and efficient indexing of feature vectors (Fig. 1a and Extended Data Fig. 1 - Stage 2). The online retrieval allows parallel image-to-image, and gene transcriptomics-to-image search functions (Fig. 1b).

**Fig 1:**
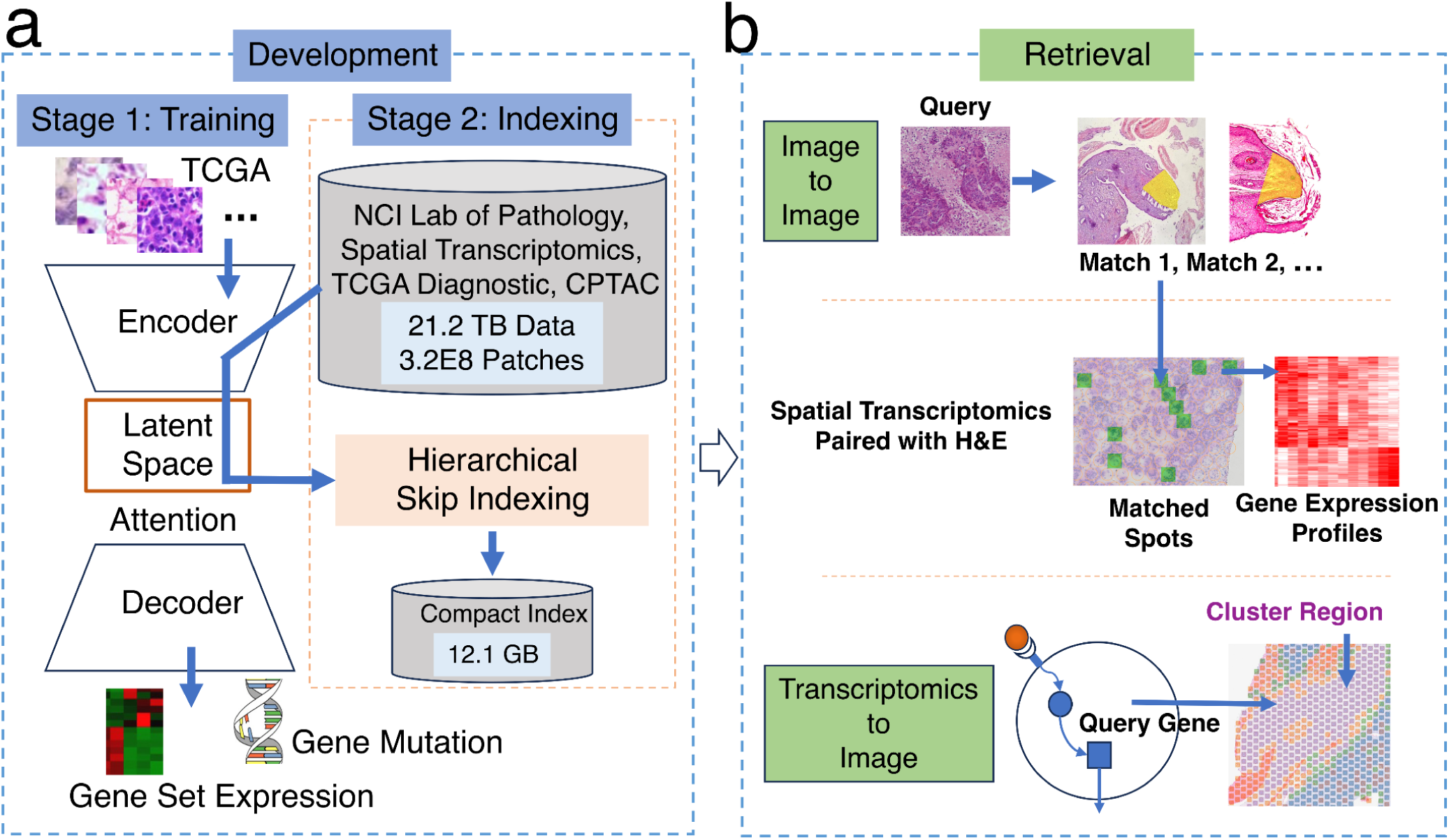
The H&E Retrieval Engine (HERE). The framework comprises two modules: Development and Retrieval. **a,** The engine development has two stages: model training and index generation. Stage 1 trains a prediction model composed of an image encoder, attention map, and decoder using TCGA data from 32 tumor types. The input to the image encoder is H&E whole-slide images, and the decoder output includes a binary indication of gene mutation functions in cancer progression and average expression of pathway gene sets. Stage 2 indexes patch vectors encoded from the whole slide images from the NCI Lab of Pathology, TCGA, CPTAC, and spatial transcriptomics studies. **b,** Two retrieval applications. For a query H&E image patch or region, the Image-to-Image retrieval returns similar image patches and associated whole slides from the indexed database. If users select images with spatial transcriptomics (ST) data, HERE returns images that closely match the query and the corresponding gene expression profiles from ST detection spots. The Transcriptomics-to-Image retrieval finds H&E slides with paired ST data where the query gene has high expression levels in ST detection spots associated with specific image feature clusters from those slides.

### Image encoder selection by tissue classification performance

The first critical step of image retrieval is encoding an input image into a numerical vector with unified dimensionality, which enables similarity comparisons with vectors encoded from database images. An input image may have various sizes, so current AI frameworks typically split the whole image into patches of a fixed size, encode patches into vectors with unified dimensionality, and then summarize all patch vectors into one vector for the entire image through attention maps^31^. Thus, stage 1 of the HERE development module establishes an image patch encoder and attention map for later retrieval functions (Fig. 1a).

A reliable H&E patch encoder should accurately capture histopathological differences, such as cell morphologies and tissue structures (Extended Data Fig. 1 - Stage 1: Tissue classification). We compared ten image patch encoders, including three neural networks trained on natural images (MobileNetV3^32^, CLIP^33^, DenseNet^34^) and seven neural networks tuned on H&E images (ProvGigaPath^35^, UNI^36^, CONCH^37^, PLIP^27^, HIPT^38^, KimiaNet^20^, and RetCCL^23^) (Extended Data Fig. 2). For the comparison, we utilized standard evaluation datasets, which are annotated H&E images from human clinical samples, including the Kather100K^39^ tissue class annotations on colorectal cancer and normal tissues, the PanNuke^40^ nuclei type annotations from 19 organ types, the BCSS^41^ region type labels for breast tumors, and the NuCLS^42^ nuclei labels from breast tumors.

Among all encoders, UNI, ProvGigaPath, CONCH, and PLIP are the top four performers with respect to differentiating image patches within the same annotation class from patches from distinct classes (Extended Data Fig. 2a, b). Although UNI and ProvGigaPath rank high in performance, their computational cost, quantified as the floating-point operation (FLOP), is over three times greater than any other encoder (Extended Data Fig. 2c). We therefore further evaluated the top four encoders for additional performance metrics relative to computational costs, which is particularly essential for the retrieval engine to avoid lengthy processing time for large input images.

### Image encoder selection by molecular prediction performance

For each top patch encoder backbone (Extended Data Fig. 2a), we trained a prediction model that encodes a whole H&E image input into a vector and predicts tumor molecular status (Methods and Extended Data Fig. 1 - Stage 1: Molecular prediction)^43^. This multi-task model can predict known effects of gene mutations on tumor progression^44^, the average expression of gene sets related to biological pathways^45^, and gene signatures related to antitumor immune response^46^ (Extended Data Fig. 3a and Methods). The training data is from TCGA, which provided three data types for individual tumor samples: diagnostic H&E slides, somatic mutations, and gene expression profiles. The collection of 6909 samples spanned 32 tumor types. We utilized each pre-trained encoder to generate image patch vectors and then added one network layer to compress the encoded vector size to 256 dimensions. Finally, we employed a multi-patch attention map (example in Extended Data Fig. 3b) to derive the case-level vector representation for each slide. The case-level feature vector can predict gene mutation and pathway expression information via the prediction network layers.

Trained with the train/validation/test scheme (Methods), CONCH, UNI, and ProvGigaPath achieved the top performance in test data (Extended Data Fig. 3c-e). We summarized performance metrics of image-patch classification (Extended Data Fig. 2a), molecular status prediction (test metrics in Extended Data Fig. 3c), and the computational cost (Extended Data Fig. 2c) in one plot (Extended Data Fig. 3f). CONCH achieved the best overall performance and maintained an over 3-fold lower computational cost than UNI and ProvGigaPath (Extended Data Fig. 3f), so we selected CONCH as the HERE backbone.

The HERE encoder comprises the pre-trained CONCH backbone plus one fully connected layer trained together with attention maps and prediction layers to predict tumor molecular status. We confirmed that the inclusion of the attention map in our framework enhanced performance relative to a framework without the attention map (Extended Data Fig. 3g). The HERE encoder converts each H&E patch with 256 x 256 pixels at 20X resolution (Methods) into a vector with 256 dimensions, a trade-off between embedding size and encoding quality. Compared to its CONCH backbone, the HERE encoder has the additional capability of predicting gene mutations and pathway expressions.

In the retrieval process, if the input image size covers multiple patches, HERE encodes the image into one vector using the patch encoder and attention map. HERE then computes the similarities between the input vector and candidate vectors encoded from image patches in the database. HERE outputs the top similar image patches with locations marked in the associated whole slide images, patient clinical information (if available), and tumor molecular status (mutations and pathways) predicted from the vector encoded from the query image.

### Limitations of the molecular prediction framework

To further evaluate prediction performance, we utilized a completely independent dataset, the Clinical Proteomic Tumor Analysis Consortium (CPTAC)^47^ cohort data. Like the TCGA training data, CPTAC contains tumor H&E images and associated bulk transcriptomics and mutation profiles. Performance metrics using the CPTAC dataset are statistically better than random expectation, defined as Area Under the receiver operating characteristic Curves (AUC) = 0.5, although the overall metrics are lower than those achieved with the TCGA held-out test data (Extended Data Fig. 3h, i). For TCGA, 6 out of 10 genes remained meaningfully better than random (AUC > 0.8, Extended Data Fig. 3h). However, none achieved meaningful performance (AUC > 0.8) in CPTAC cohorts despite the statistical significance compared to random expectations (AUC = 0.5).

This performance deterioration of the TCGA-trained predictor on the CPTAC test data reflects a common challenge: Most AI predictors deteriorate when applied to test data generated in a setting different from that in which their training data are generated^48^. Constructing a reliable predictor requires training data from similar cohorts, collected under the same protocol and with similar patient and disease compositions^49^.

We further analyzed a few example predictions on the *TCGA test set*, where the training and testing cohorts are matched, and overall prediction performance remains high (Extended Data Fig. 3h). This setting allows us to dissect failure modes that arise even under favorable conditions. In a representative *correct prediction case* (Extended Data Fig. 4a), all three mutations were accurately predicted, and the attention map highlighted patches from tumor regions with consistent histological patterns. In contrast, in a representative *failure case* related to an endometrial tumor (Extended Data Fig. 4b), all four mutations were incorrectly predicted as negative. The corresponding attention map selected patches with misleading or heterogeneous features. Specifically, *cluster 0* displayed squamous differentiation—regions where cancer cells resemble squamous rather than typical adenocarcinoma morphology of endometrial tumors. Clusters 1, 2, and 6 contained high vessel density, while clusters 3, 5, and 7 primarily showed red blood cells—features that are not tumor cells. These non-representative patches, despite receiving high attention scores, may have contributed to the incorrect predictions by deviating from canonical cancer cell morphologies learned during training.

This analysis underscores a key limitation: the model can be misled by non-canonical histological patterns, particularly in cases with heterogeneous or atypical tumor features. Carefully selected training data with sufficient corner case coverage will be necessary to improve AI performance further.

### Indexing large numbers of high-dimensional vectors

An image encoder cannot be directly utilized as a retrieval engine because many additional functions, such as database indexing (Extended Data Fig. 1 - Stage 2) and result prioritization, still need to be implemented. The biggest challenge in developing an H&E retrieval engine is finding similarities across a large number of vectors in a way that is efficient with time and memory. We collected about 21.2 terabytes of H&E images of various human tumor types from the NCI Lab of Pathology (11232 whole-slide images, 7.12 terabytes), TCGA project (11738 whole-slide images, 11.78 terabytes), CPTAC (7335 whole-slide images, 2.22 terabytes), and spatial transcriptomics studies (130 whole-slide images, 87.4 gigabytes). Encoding image patches into vectors using the patch encoder trained as above produced about 3.2E8 vectors in 256 dimensions. Brute force comparisons between an input vector and all candidates would demand tremendous computation time and memory space.

Evaluation of possible solutions revealed that the freely available Facebook AI Similarity Search (FAISS) library^50^ provides reliable and rapid large-scale comparisons of H&E vectors. The FAISS library provides high-dimensional vector indexing to accelerate nearest-neighbor searches at a tradeoff of accuracy loss. We evaluated two FAISS solutions. One named HNSW-IVFPQ combines three algorithm designs, including Hierarchical Navigable Small World (HNSW)^51^, Inverted File Index (IVF)^52^, and Product Quantization (PQ)^53^ (Fig. 2a). The PQ algorithm reduces the size of high-dimensional vectors while minimizing information loss^53^. For example, all of the vectors for the NCI, CPTAC, and TCGA datasets described above can be loaded into 12.1 gigabytes of random-access memory, demonstrating that all vectors for even very large databases can be loaded into the memory of a personal computer. The IVF algorithm confines the search scope to a subset of vectors around a centroid vector close to the query input^52^, while the HNSW algorithm efficiently locates the IVF centroid through hierarchical skip indexing^51^. The second FAISS solution is ITQ-LSH, which combines Iterative Quantization (ITQ)^54^ and Locality Sensitive Hashing (LSH)^55^ algorithms to convert high-dimensional vectors into compact binary codes.

**Fig 2:**
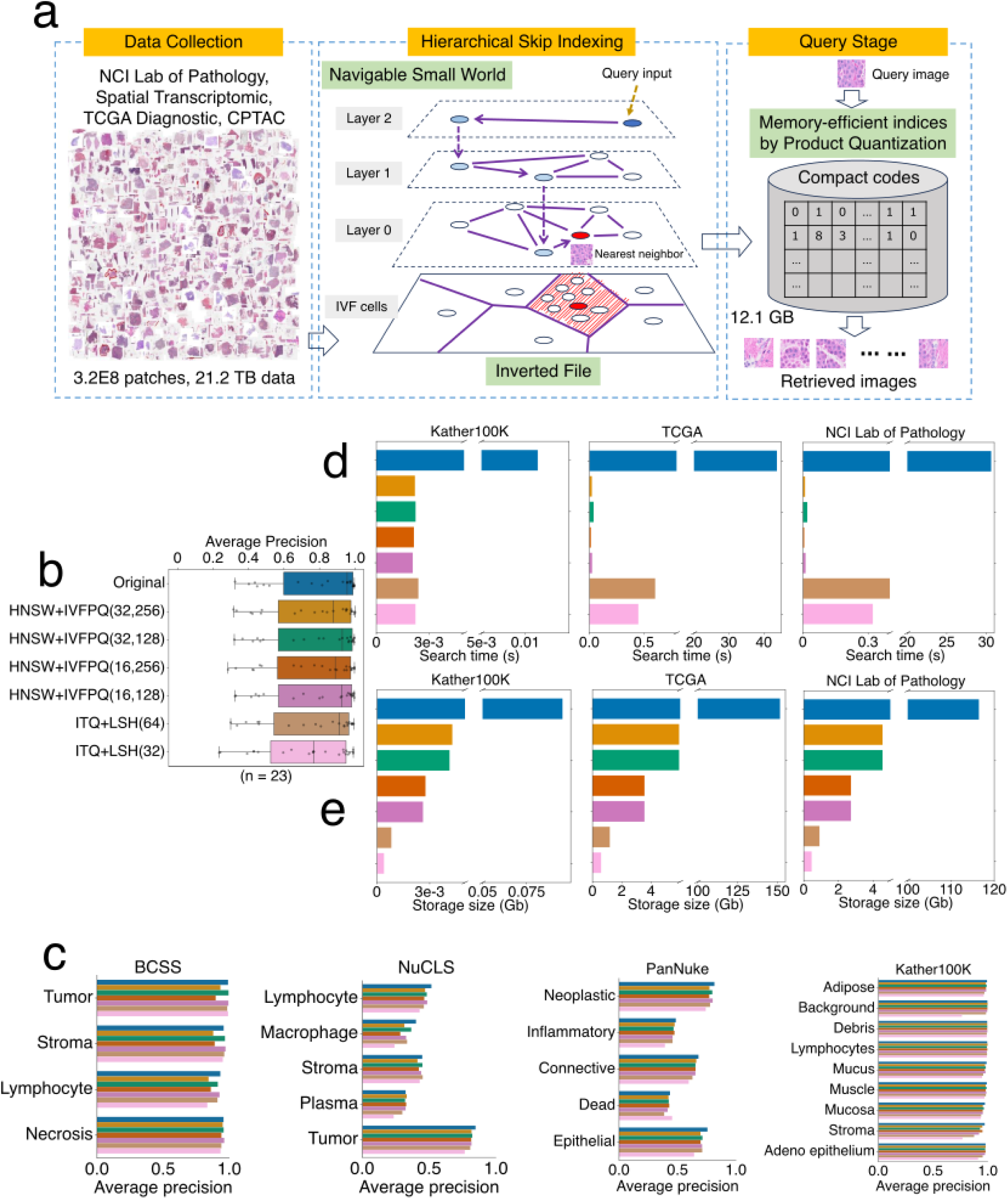
Indexing large numbers of feature vectors. **a**, Hierarchical skip indexing using the FAISS library. The Product Quantization (PQ) algorithm reduces dense vectors into memory-compact code. For a query vector, the Hierarchical Navigable Small World (HNSW) algorithm can rapidly identify a nearby centroid, a summary point of many similar vectors. Then, the Inverted File (IVF) Index lists a subset of candidates around the centroid for searching. **b**, Accuracy comparisons among vector indexing approaches. The FAISS library provides two representative vector indexing approaches, HNSW-IVF-PQ (panel a) and ITQ-LSH (Methods). Each indexing approach has different parameter settings indicated in the brackets (Methods). The average precisions across all classes and datasets (details in panel c) were compared between each method and brute-force searching with original vectors. The thick line represents the median value. The bottom and top of the boxes are the 25th and 75th percentiles, respectively (interquartile range). Whiskers encompass 1.5 times the interquartile range. **c**, Accuracy comparisons among vector indexing approaches in individual datasets. Four H&E annotation datasets, including BCSS, NuCLS, PanNuke, and Kather100K, were used to evaluate the average precision (across all patches) of top-5 results from different vector indexing approaches. **d**, Time comparisons. We randomly selected 100 image patches from the BCSS dataset as query inputs and queried top-10 results from each cohort. The average time among 100 queries was shown. **e**, Index size comparisons for each cohort.

We compared the image retrieval performance of these two FAISS solutions (ITQ-LSH and HNSW-IVFPQ) with a brute-force search using the original high-dimensional vectors. HNSW-IVFPQ achieved the best overall retrieval performance on four benchmark datasets, with a negligible accuracy decrease compared to the brute-force search using original vectors (Fig. 2b,c).

HNSW-IVFPQ substantially improved retrieval speed and memory consumption (Fig. 2d, e). Using compact indexing by HNSW-IVFPQ across evaluation standard datasets, the average time for retrieving top-10 similar items is less than 0.1 second, compared to about 30 seconds by brute-force search using the original features (Fig. 2d). Likewise, the original dense vectors generated with NCI Lab of Pathology data occupied 116 gigabytes of memory, while the compact 8-byte codes generated by HNSW-IVFPQ occupied about 4.6 gigabytes (Fig. 2e). Further scalability analyses revealed that the indexing strategy can enable retrievals within seconds and contain the database index size within 16GB (the typical memory of a desktop computer) even for the combined set of images from the TCGA, CPTAC, and NCI Lab of Pathology (Extended Data Fig. 5).

### Image-to-Image search module on the HERE web app

With the encoder and database index above, we deployed a public web portal for large-scale H&E image retrieval. Users upload one image region of interest to retrieve similar image patches from the H&E collection from the NCI Lab of Pathology, TCGA, or CPTAC projects. HERE encodes the input image into a vector with the encoder and attention map described in previous sections. Then, the system will return top-K (50 by default) most similar image patches and their locations in the corresponding whole-tumor slides, together with clinical information if available. The result page displays whole-slide images ordered by the sum of rank scores from the top-K retrievals. HERE also computes the statistical significance (empirical z-score and p-value) for each top patch by comparing the distance between the input vector and top patch vector against background distances, which were calculated between the input vector and 10,000 randomly selected patches.

### Blinded pathologist evaluation using the HERE101 benchmark

To assess inter-observer variability and evaluate the reliability of the HERE web portal, we performed a blinded evaluation with three human pathologists (Extended Data Fig. 1, Stage 3). First, a pathologist designed a benchmark called “HERE101” that consists of 101 diagnostic H&E images from different human tumors and organs (Fig. 3). The benchmark covers six characteristics to describe tumors. Tumor/normal content (aspect 1) indicates whether the current image is from tumor or normal tissues. Cytoplasmic features (aspect 2) include dichroic, basophilic, eosinophilic, clear, and scant. The tissue site (aspect 3) indicates the organ locations. Tissue structures (aspect 4) include cord, nest, cribriform, sheet, microcystic, swirl, glandular, trabecular, bundle, stratified, and papillary.

**Fig 3:**
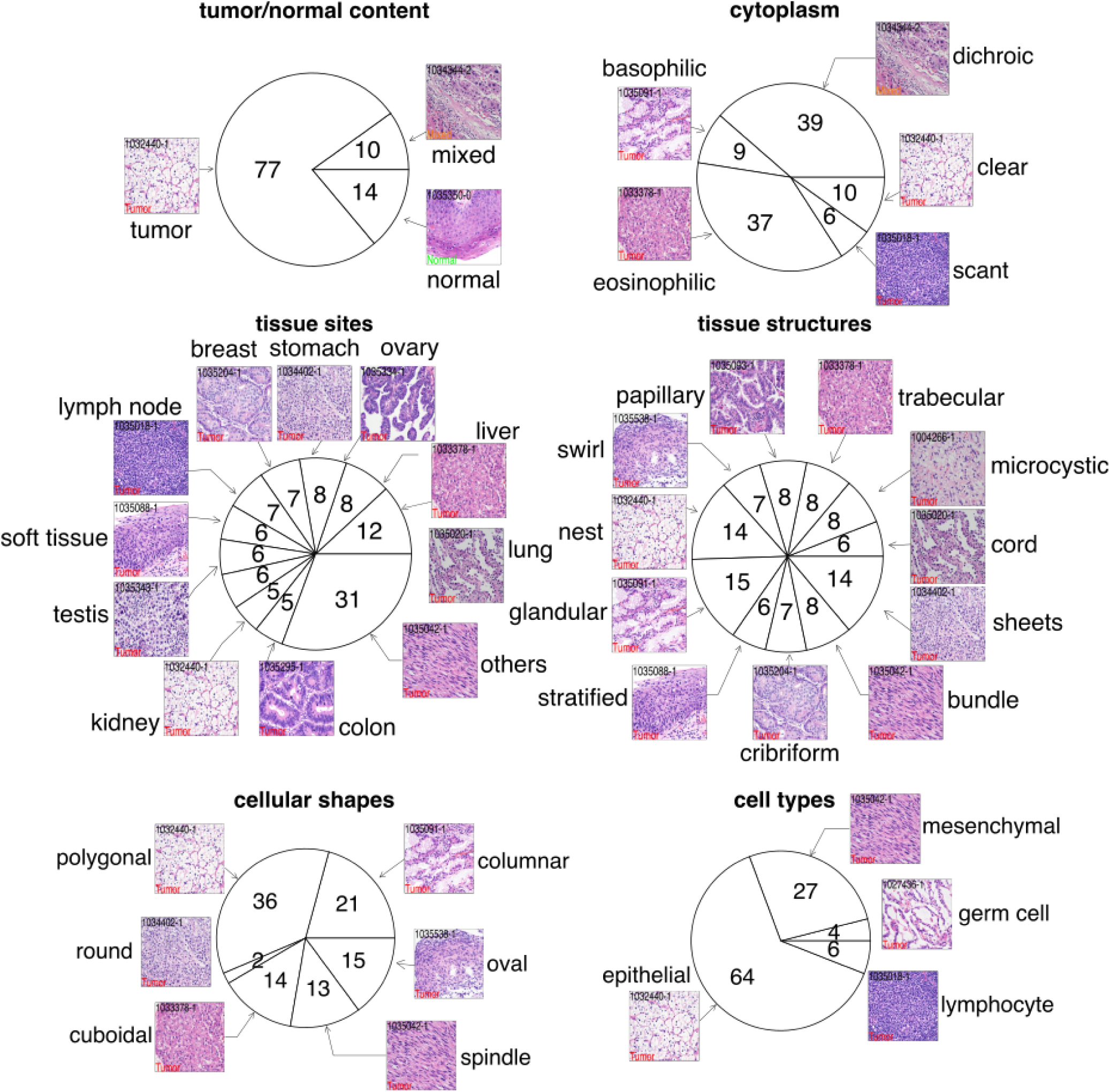
The HERE101 benchmark for evaluating H&E retrieval performance. The HERE101 benchmark includes 101 diagnostic H&E images from different human tissues. The pathologist selected images to represent the diversity of six categories of image features. For each group, the numbers in each pie chart slice indicate the number of cases. One representative image is shown for each slice.

Cell shapes (aspect 5) include polygonal, columnar, cuboidal, spindle, round, and oval. Cell types (aspect 6) include epithelial cells, lymphocytes, mesenchymal cells (e.g., smooth muscle cells, adipose cells, Schwann cells, endothelial cells, etc.), and germ cells.

Three pathologists used each image in HERE101 as a query to return the top 50 similar patches and associated whole slides from the NCI Lab of Pathology collection (Fig. 4a). We selected the NCI in-house data for the HERE101 test for two reasons. First, we used TCGA data to train the encoder. By using NCI data, we avoided using TCGA data again in the validation. Second, we hope hospitals and clinical centers widely use the HERE framework in searching their in-house records. Using the NCI in-house data is a practical proof-of-concept demonstration for using the HERE search engine in the context of a specific clinical center.

**Fig 4:**
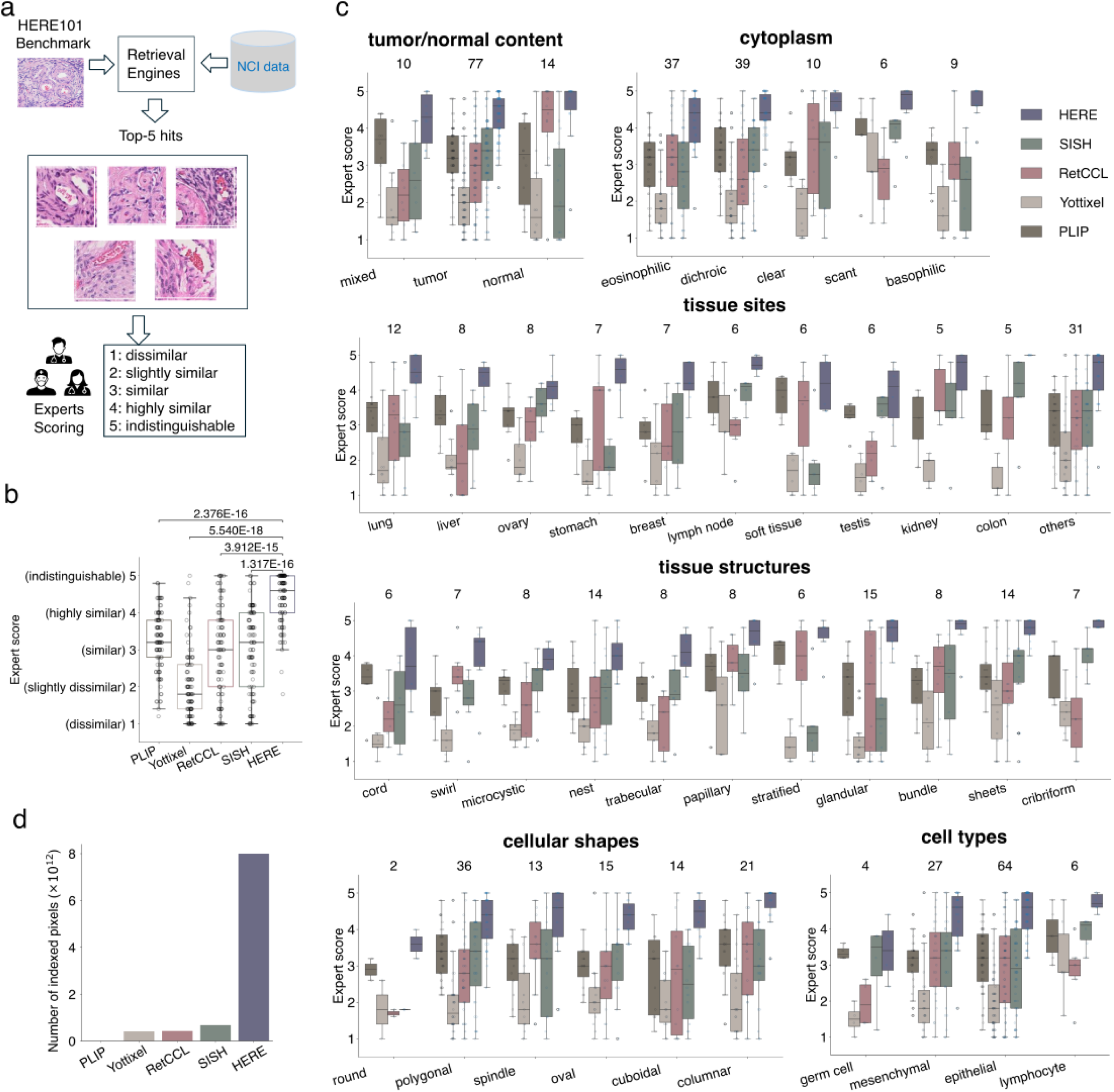
Retrieval accuracy assessment on HERE101 benchmark. **a**, The blind evaluation procedure with three clinical pathologists. For each method, we submitted each of the 101 images in the HERE101 benchmark (details in Fig. 3) and scored the top 5 whole-slide images returned from searching the NCI Lab of Pathology data. **b**, Performance scores on the HERE101 benchmark evaluated by three pathologists. Within the box plot, each dot represents the median scores from three pathologists among the top 5 whole-slide images for a query (n= 101 per box group). The thick line represents the median value. The bottom and top of the boxes are the 25th and 75th percentiles, respectively (interquartile range). Whiskers encompass 1.5 times the interquartile range. The P-value was calculated using the two-sided Wilcoxon signed-rank test, comparing scores between HERE and other methods paired on each input image. The performance scores from individual pathologist are summarized in Extended Data Fig. 7. **c**, Performance scores are grouped by image features and shown as panel b, with the number of cases per group labeled on the top. **d**, Total image area (by pixels) included for retrieval by each method on the NCI Lab of Pathology data (SISH, RetCCL, Yottixel) or default database provided by the web portal (PLIP).

Each pathologist then visually examined the top five whole-slide images and scored each whole slide on a five-point whole-number scale, based on the similarity to the input image: 1 (dissimilar), 2 (slightly similar), 3 (similar), 4 (highly similar), 5 (clinically indistinguishable) (Fig. 4a and Extended Data Fig. 6). We compared HERE with previously published CBIR frameworks, including SISH^21^, Yottixel^19^, and RetCCL^23^, all of which provide source code (but no web portal), along with the PLIP web portal (WebPLIP)^27^. SISH^21^, Yottixel^19^, and RetCCL^23^ extract a small fraction of patches from each whole-slide image to reduce the search space. We ran the source code of each framework with default parameters on the NCI Lab of Pathology data for the HERE101 benchmark retrievals. PLIP^27^ reduces the entire image into one patch for encoding, thus cannot preserve resolution information of whole-slide images from the NCI data. Therefore, we utilized the PLIP’s pre-processed validation images as the database for the HERE101 benchmark retrievals. To ensure blindness, we masked the framework names and unified the output configurations (Methods); thus, pathologists cannot identify the method identities during the scoring.

Compared to other CBIR frameworks, HERE achieved significantly higher scores based on the consensus of pathologists (Fig. 4b and Extended Data Fig. 7a). 12.9% of HERE retrievals achieved a score of 5 (clinically indistinguishable from input) and 45.5% scored 4 (highly similar) (Fig. 4b), meaning 58.4% of retrieved slides were “highly similar” or better. Furthermore, on each of the six image characteristics, HERE achieved the best performance among all methods on all categories (Fig. 4c). On the NCI Lab of Pathology data, the image area indexed by HERE is 19-fold greater than Yottixel, 18.5-fold greater than RetCCL, and 12-fold greater than SISH (Fig. 4d).

To assess the inter-rater agreement and consistency among the three pathologists, we computed the pairwise Pearson correlations between ratings across all cases and methods (Extended Data Fig. 7b). Most of the Pearson correlations between pathologists’ ratings ranged from 0.5 to 0.75, suggesting a moderate level of inter-rater agreement.

### Performance evaluation on cancer type and mutation matching

To further compare retrieval frameworks (Extended Data Fig. 1, Stage 3), we utilized data from the Clinical Proteomic Tumor Analysis Consortium^47^ (CPTAC), a cohort similar to TCGA with a matched set of tumor H&E images, bulk transcriptomics, and mutation profiles. We did not utilize CPTAC for any previous training and testing procedures, so CPTAC could serve as an independent test set.

First, we tested whether using a whole-slide image (WSI) to query CPTAC with each framework returns WSIs from the same cancer type among the rest of the CPTAC images. Among all methods, HERE achieved the best overall average prediction (Fig. 5a) and the top performance in 12 out 13 total cancer types (Fig. 5b). Second, we tested whether using a WSI as a query returns top hits from tumors with the same mutation status. HERE achieved the best overall performance (Fig. 5c) and the top performance in 8 out of 10 high-frequency mutations (Fig. 5d). Together, these results further support the superior performance of HERE relative to existing retrieval frameworks.

**Fig 5:**
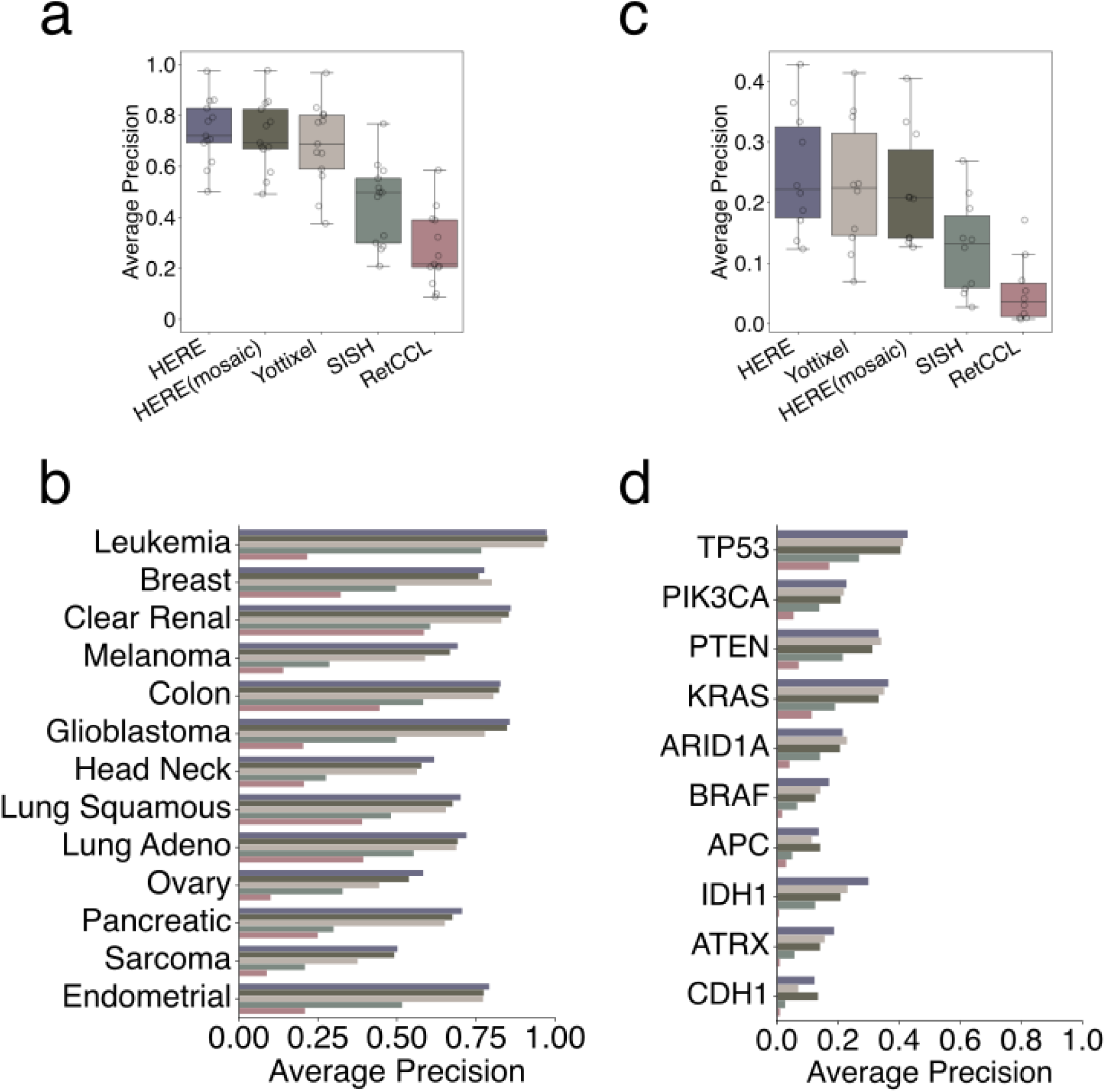
Retrieval accuracy comparisons based on case-matching in omics cohorts. To further compare retrieval frameworks, we tested the consistency of cancer types (a, b) and mutations (c, d) among the top five images returned. The test data come from CPTAC cohorts, which were not utilized in any training steps. We utilized a leave-one-slide-out validation procedure. For each whole-slide image (WSI) as input, we used the Yottixel mosaic selection procedure to identify the most informative patch. Then, for each query patch selected for a WSI, we retrieved the top 5 WSIs from the remaining WSIs in the whole cohort and computed the mean average precision^21^ (Methods) using the ground-truth label of the input image. **c**, Summary of average precision in matching tumors with the same mutations. Each dot represents a mutation from the top 10 most frequent mutations that are included in both TCGA and CPTAC (details in panel d). The result is shown in box plots as Fig. 4b. **d**, Average precision in matching tumors with the same mutations.

In this performance analysis, Yottixel achieved close performance to HERE (Fig. 5a). Unlike HERE that indexed all image patches, Yottixel utilized a mosaic approach to sub-sample image patches to reduce the search space. The close performance indicates that the mosaic procedure in Yottixel may enhance the scalability of HERE by reducing the search space to the most representative patches from whole slide images. Using CPTAC data, we tested the HERE(mosaic), indexing the mosaic-selected (20%) patches from each image (Methods). For both cancer type and mutation retrieval tasks, HERE(mosaic) achieves performance comparable to the original HERE (Fig. 5). Meanwhile, the mosaic strategy reduced the HERE index on the CPTAC data from 1.6 to 0.27 gigabytes (i.e., 6-fold reduction) in the random access memory. This result suggests that the mosaic selection can enhance the scalability of the HERE framework without significantly undermining the retrieval performance.

### Inference of expressed genes in a query image

H&E images alone do not provide any genetic or molecular information. Spatial transcriptomics (ST) bridges image and molecular information, allowing concurrent profiling of H&E images and gene expression on the same tissue slide^56^. Searching H&E images with ST data obtained on the same slide can provide users with gene expression profiles from ST detection spots in the matched image patches (Fig. 1b). HERE contains 130 ST profiles paired with H&E images from public studies (Supplementary Table 1).

To illustrate this capability, for an image query, we used an H&E image from a tumor hypoxia region marked by anti-Pimonidazole staining^57^. We restricted retrievals to H&E images accompanied by ST data from the same slide. HERE returned 50 H&E-ST image patches from 4 different tumor types (colorectal, lung, ovarian, and breast; example patches in Extended Data Fig. 8a). These ST profiles covered 18,031 human genes, each of which was detected in at least 75% of ST spots in the returned image patches (example genes in Extended Data Fig. 8b). Gene set enrichment analysis^58^ on the mean expression across all spots identified seven pathways (q-value < 0.05 and enrichment scores higher than hypoxia) with glycolysis as a top term (Extended Data Fig. 8c, d). Previous studies show that tumor hypoxia (the lack of oxygen) activates genes involved in glycolysis (free energy generation without oxygen)^59^.

The top two genes by mean expression across matched ST spots are *VEGFA* and *SLC2A1* (Extended Data Fig. 8b). VEGFA is a well-known tumor hypoxia marker and a cancer therapeutic target^59^. SLC2A1, also named GLUT1, is a membrane protein facilitating glucose uptake in most cell types, including cancer cells^60^. Thus, HERE can find clinically relevant biomarkers from image retrievals.

### Transcriptomics-to-Image search module

Spatial transcriptomics (ST) data paired with H&E images also makes it possible to implement a Transcriptomics-to-Image module, which returns image features associated with the high expression level of a query gene name (e.g., *SERPING1*) (Fig. 1b). A simple approach would be to return image patches in which ST data shows high expression of a query gene. However, such image patches would be created without consideration of image similarities among patches, precluding meaningful generalization of image features associated with the gene expression levels.

To address this hurdle, we pre-calculated statistical links between gene expression and image features, defined as clusters of image patches with similar encoding vectors. First, for each ST profile, we clustered image patches around ST detection spots into eight clusters based on cosine similarities (equivalent to Euclidean distances in our implementation, Methods) between the image-patch encoding vectors using the k-means algorithm. We conducted the k-mean clustering within each ST profile but not across all ST profiles because we observed batch effects, reflected as patches from the same sample clustering together (Extended Data Fig. 9a). Then, within each ST profile (from the same parent image), we tested whether a query gene’s expression over detection spots within the image cluster is significantly higher than spots outside the cluster (Fig. 6a). For each image cluster, HERE generates a vector of Cohen’s d values and rank-sum test p-values for all profiled genes, quantifying the association between gene expression and image features represented by that cluster (Fig. 6a bottom box).

**Fig 6:**
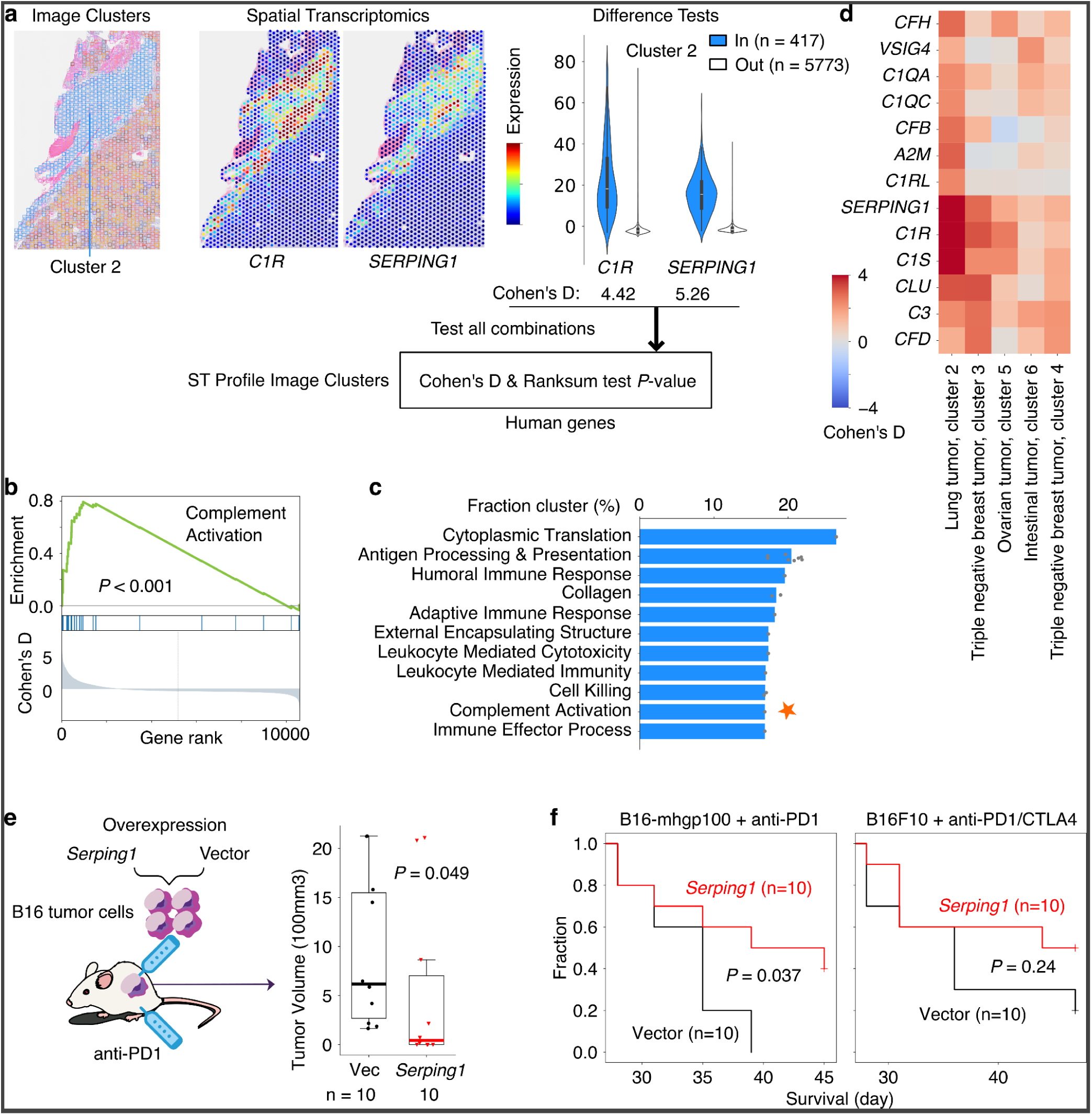
Transcriptomics-to-Image search: HERE returns image features given an input gene. **a**, Associations between gene expression and image features. Hierarchical clustering was applied to Spatial transcriptomics (ST) data from a human lung tumor to organize image patches around ST detection spots into eight clusters (left panel; each cluster is a different color). For a given query gene (e.g., *C1R* and *SERPING1,* center panel) and each image cluster (for example, cluster #2 (blue) in left panel), the expression difference among ST detection spots within the image cluster region and spots outside the cluster region is quantified using the Cohen’s d value (right panel). Testing each of all possible query genes against each of every ST profile cluster will generate the result matrix (bottom panel). **b**, Complement Activation gene set enrichment. Along the x-axis, all genes are ranked from high to low by Cohen’s d values (bottom Y-axis) computed for cluster #2 of the ST profile in panel a. Members of the “complement activation” pathway from Gene Ontology biological processes (GO_BP)^45^ are indicated by horizontal blue lines in the middle of the plot. The top y-axis plots “complement activation” enrichment scores at each gene rank. The *P*-value is computed through the one-sided permutation test (1000 randomizations). **c**, Top 20 GO_BP terms associated with image features. For each term, the Y-axis presents the fraction of ST profile clusters whose Cohen’s d gene scores are significantly enriched (False Discovery Rate < 0.05). Multiple GO_BP terms related to similar biological processes are grouped with the y-axis presenting mean values across all merged terms. The star marks the complement activation pathway discussed in the main text. **d**, Cohen’s d heatmap of complement activation genes. Columns are labeled with each ST profile’s cancer type and the cluster index. **e**, In-vivo effects of *Serping1* overexpression on tumor volume. Left panel: B16-mhgp100 cells with *Serping1* and vector-only overexpression were inoculated subcutaneously into mice treated by immune checkpoint blockade. Right panel: The tumor sizes on day 28, the day before the first tumor reached an endpoint (tumor volume ≥ 2000 mm^3^ or length ≥ 2 cm). Box plots are shown as in Fig. 4b. Group values were compared through the two-sided Wilcoxon rank-sum test. **f**, *Serping1* overexpression in tumors extended survival. B16-mhgp100 cells express immunogenic antigen hgp100, while the B16F10 cell line is less immunogenic. On the Y-axis, the fraction of mice with endpoint-free survival is plotted against days since tumor inoculation (X-axis). The two-sided log-rank test compared group survival differences.

Given a query gene, HERE returns all image clusters with a significantly high level of query gene expression relative to other clusters on the parent slide (Cohen’s d > 1 and false-discovery rate (FDR) < 0.05, Methods). On average, across all slides, each cluster of ST image patches has 3999 genes with significant associations (FDR < 0.05, Extended Data Fig. 9b). This indicates that ST image patch clusters, which represent image features, also captured characteristic gene expression patterns within tumors.

### Molecular pathways associated with tumor morphologies

The pre-calculated Cohen’s d values relating gene expression and image features not only enable the transcriptomics-to-image search module but also provide an opportunity to reveal genes and pathways associated with tumor morphologies systematically (Extended Data Fig. 1, Stage 4). We performed gene set enrichment on the Cohen’s d profiles (Fig. 6b) across all 1039 image clusters and found 783 pathways (of 3211) that were significantly enriched in at least one cluster (FDR < 0.05, Extended Data Fig. 9c). We ranked the enriched pathways according to the fraction of ST clusters in which each pathway is enriched. The commonly enriched pathways (Fig. 6c) involve adaptive immune response and other processes associated with tumor histologic morphology. For example, antigen processing & presentation is a hallmark of tumor-infiltrating T cells, which have distinctive nuclear morphology in H&E images^61^, while extracellular collagen contributes to the fibrotic morphology in tumor stroma^62^.

Most top enriched pathways (Fig. 6c) have well-defined roles in cancer (Supplementary Table 2). One notable exception is complement activation, an innate immunity pathway typically activated by pathogens or pathogen-bound antibodies, resulting in a cascade of plasma protein reactions that induce inflammatory responses^63^. The complement system has a traditional role in fighting infectious diseases. However, more recent studies have shown that complement activation can act as either a negative or positive regulator of tumorigenesis^64^. Prompted by the HERE results and the emerging evidence for the role of the complement system in cancer, we investigated further.

We noted that genes enriched in our analysis come mainly from the complement component 1q (C1Q) complex, including *C1QA/B/C*, *C1R*/*S*, and Serpin Family G Member 1 (*SERPING1*) (Fig. 6d, one-sided hypergeometric *p*-value = 6.6e-4). These C1Q genes are expressed in tumor fibrotic stroma of the top-matched ST images (Fig. 6a and Extended Data Fig. 9d). Mice deficient in C1Q display decreased tumor growth^65^, suggesting that activation of the C1Q promotes tumor growth. Additional evidence comes from the knockdown of C1Q components C1R and C1S in cancer cell lines, which slows xenograft tumor growth in mice^66^. In the classical activation pathway, the C1Q recognition proteins interact with proteases that cleave downstream proteins in the cascade. SERPING1 inhibits C1Q activation by binding to the C1R/S protease, causing it to dissociate from C1Q^63^.

Given the pro-tumor roles of the C1Q complex, we hypothesized that the C1Q inhibitor SERPING1 may inhibit tumor progression. As an initial investigation, we overexpressed the C1 inhibitor gene *Serping1* in B16 tumor cell lines and measured the B16 tumor growth under immune checkpoint blockade (Fig. 6e). RT-qPCR analysis confirmed a 552.6-fold change in *Serping1* expression relative to the empty vector. Compared to the vector control, mouse *Serping1* overexpression significantly (*p* = 0.037) slowed in-vivo growth of the B16-mhgp100 cell line, a syngeneic model that expresses immunogenic antigen hgp100^67^ (Fig. 6e, f). In the B16F10 cell line, which is less immunogenic, the *Serping1* overexpression also slowed tumor growth (Fig. 6f), albeit not to a level of statistical significance (*p* = 0.214). We did not observe any changes in B16F10 cell line growth *in vitro* upon *Serping1* overexpression (Extended Data Fig. 9e), showing that an immune response is required for the inhibitory effect. These results suggest that the complement system may modulate tumor progression under immune checkpoint blockade.

## Discussion

We developed the HERE framework, which features Image-to-Image and Transcriptomics-to-Image searches with large-scale H&E image databases. Currently, HERE utilizes the NCI Lab of Pathology, TCGA, and CPTAC databases, and features a web portal that provides public access. Unlike previous approaches that downsample whole-slide images, HERE scans all regions of candidate database images at high resolution. The 21.2 terabytes of H&E images in the NCI, TCGA, and CPTAC databases were used to build an index of 12.1 gigabytes, which can be loaded entirely into the memory of a desktop computer with a 16-gigabyte memory.

Many recent studies in digital pathology AI aim to build foundational models on unified feature extraction for different tasks, such as clinical label or outcome predictions. We emphasize that retrieval engines and prediction models have distinct purposes and thus are complementary. For example, clinicians may want to use a retrieval engine to search for similar cases in the same hospital (e.g., the NCI Lab of Pathology cohort) or from public studies (e.g., TCGA or CPTAC cohorts), which may provide associated information such as previous diagnosis and genomics profiles for example-based interpretability.

To evaluate HERE, we designed and utilized a benchmark standard called HERE101, which consists of 101 images that cover the wide range of cell types, tissue sites, structures, cell shapes, and cytoplasm features found in tumor H&E images. Pathologist evaluation of web portals using HERE101 demonstrated improved performance of HERE relative to previous retrieval tools. The HERE101 benchmark may serve as a general standard for evaluating future H&E retrieval web portals.

The HERE web portal enables users to search and explore whole-slide image (WSI) collections from the NCI Pathology Lab and NCI-sponsored projects, including TCGA and CPTAC. Currently, the portal does not support user-uploaded data due to the substantial computational and storage demands of hosting high-resolution WSIs, encoding new images, and updating the retrieval index in real time. For users who wish to query their own datasets, we provide open-source code and detailed instructions for local deployment. While the current implementation does not support community uploads, the HERE infrastructure is designed to allow incremental updates to the image index. As such, enabling user-upload functionality remains a feasible direction contingent on future computational and budgetary resources.

HERE bridges the gap between gene molecular information and image features. The training of the image encoder aims to predict the status of gene activity underlying a query image. These include the potential effects of specific mutations and the average gene expression of pathways. The availability of spatial transcriptomics (ST) data paired with H&E images enabled HERE to link molecular biology and histopathology. The image-to-image query module can return gene expression profiles for ST spots with images that closely match the query. Further, the transcriptomics-to-image retrieval can suggest image features associated with a gene of interest. A limitation of these functions is the low sample size of 130 ST profiles, which may not sufficiently cover the associations between molecular and image features. Expanding ST datasets would be crucial for making this functionality more robust in future improvements.

Using associations (Cohen’s d) between image features and spatial transcriptomics (ST) levels, we conducted a proof-of-concept analysis to explore molecular pathways linked to tumor morphology. As a case study, we investigated a top association involving SERPING1 and complement activation in in vivo models. This analysis illustrates that HERE has the potential to go beyond image retrieval by uncovering molecular regulators of tumor progression. However, it is essential to note that the association scores (Cohen’s d) reflect only correlations between regional gene expression and image features—they do not imply causal biological regulation, which involves complex interactions among diverse cell types and molecular processes within tumors. Dedicated frameworks will be needed to identify causal regulators of tumor histopathological patterns.

The current HERE framework has several limitations. The attention map of HERE, which performs a weighted average of patch vectors, cannot model complex multi-patch spatial organizations. Capturing tissue structures or image features that span multiple patches may require new algorithm designs. Another limitation is that the input to HERE is currently unimodal (either an image or gene name). However, clinicians may want to search with a combination of image, text, and molecular test results. Future HERE extensions will allow multimodal search and access to more complete data, including electronic health records and molecular test results. Finally, the HERE image database is still on the terabyte scale, much lower than that of Google or BLAST. Once histopathology images are available on the petabyte scale and above, the current HERE database infrastructure may need to be upgraded with additional solutions, such as distributed systems and indexing of only selected content.

One consideration for users is the alignment of resolutions between the input image and database images. The HERE framework was trained on 20X magnification images (Methods), but images input by users may have different resolutions (for example, 80X or 40X), or an unknown resolution. We recommend that users start their search using the full resolution of the input image. If the initial results are unsatisfactory or have low significance scores (z-scores and p-values), the adaptive search function on the web portal enables users to search again by reducing the query resolution by half, which may improve results by better aligning the input resolution with the database resolution. We also recommend that the input image be a region of interest with constrained sizes because the HERE encoder and attention framework cannot model complicated tissue structures across multiple patches.

HERE is immediately useful to clinicians seeking to compare their case of interest with tumors documented by TCGA, CPTAC, and NCI, and to molecular biologists interested in linking the activity of genes and pathways to tumor spatial organization. The HERE training framework also enables clinical centers to establish custom local retrieval engines on a typical desktop computer. NCI will provide long-term support for the HERE server, and we plan to expand the NCI in-house cohort once more whole-slide images become available at the NCI Lab of Pathology. Although this study focuses on cancer, the HERE framework applies to any tissue type in health or disease.

## Methods

### Test data for comparing image encoders

This section corresponds to the “Stage 1 - Tissue classification” in Extended Data Fig. 1 and Extended Data Fig. 2. To compare different pre-trained image encoder backbones, we utilized the following four datasets:

1. The Kather100K colon dataset^39^ contains 100,000 non-overlapping image patches extracted from 86 H&E images of human colorectal cancer and normal tissue. All images are 224 x 224 pixels at 0.5 microns per pixel (µm/px). Tissue classes include adipose, background, debris, lymphocytes, mucus, smooth muscle, normal colon mucosa, cancer-associated stroma, and colorectal adenocarcinoma epithelium.
2. The PanNuke dataset^40^ for nuclei segmentation comprises images from 19 organ types. Each image contains five different types of nuclei: neoplastic, inflammatory, connective tissue, epithelial, and dead. The dataset includes 189,744 annotated nuclei distributed across 7,904 images, each with a resolution of 256 × 256 pixels. The images have a resolution of 0.25 µm/px.
3. The BCSS dataset^41^ comprises 151 H&E whole-slide images from FFPE breast tumors studied in the Cancer Genome Atlas (TCGA) project^68^. Crowdsourcing efforts annotated 21 region-level labels, including tumor, stroma, lymphocyte-rich regions, necrosis, blood, fat, plasma cells, nerve, etc. We removed ambiguous labels, such as *outside_roi*, *exclude*, and *other*, and kept the four most frequent tissue labels, including tumor, stroma, lymphocytic infiltrate, and necrosis or debris.
4. The NuCLS dataset^42^ contains over 220,000 labeled nuclei from breast cancer images from TCGA^68^ through a collaborative effort involving pathologists, pathology residents, and medical students. We utilized the single-rater data with annotations reviewed by pathologists to ensure accuracy and reliability. Annotation labels include lymphocyte, macrophage, nonTIL-nonMQ stromal, plasma cells, and non-mitotic tumors.

For datasets 2, 3, and 4, we generated patch-level annotations from the nuclei or region annotations as follows. We chopped non-overlapping patches in 512 x 512 pixels (0.25 µm/px) from the original images and resized patches to 256 x 256 pixels (0.5um/px). The label with the most frequent area within the patch was then assigned as the patch label for further analysis.

### Average precision for comparing image encoders

This section corresponds to the “Stage 1 - Tissue classification” in Extended Data Fig. 1 and Extended Data Fig. 2.

We compared with ten image encoders, including three networks trained on natural images (MobileNetV3^32^, CLIP^33^, DenseNet^34^) and seven networks tuned on pathological images (ProvGigaPath^35^, UNI^36^, CONCH^37^, PLIP^27^, HIPT^38^, KimiaNet^20^, and RetCCL^23^). For each patch in a test dataset (i.e., BCSS, NuCLS, PanNuke, and Kather100K introduced above), we applied each encoder to rank the top-five closest patches by cosine similarities (equivalent to Euclidean distances in our implementation, Methods) from the remaining patches in the dataset. For each class, we report the mean average precision at 5 (mAP@5) performance^21^ defined as follows:

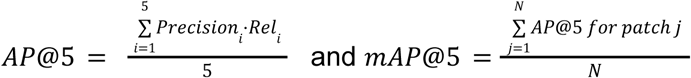

N represents the number of patches in each class. 𝑃𝑟𝑒𝑐𝑖𝑠𝑖𝑜𝑛_*i*_ represents the fraction of correct matches among cases up to i. 𝑅𝑒𝑙_*i*_ represents the correctness flag for the top 𝑖-th match. Finally, for method comparisons, the median mAP@5 across all classes and datasets is utilized (Extended Data Fig. 2a).

### Training data for the encoder and attention map

This section corresponds to the “Stage 1 - Molecular prediction” in Extended Data Fig. 1, the “Stage 1: Training” in Fig. 1a, and the multi-task neural network model (Extended Data Fig. 3a). A summary of the multimodal data availability is presented in the Supplementary Table 3.

#### 1 Image data

We collected 6909 H&E diagnostic whole-slide images with corresponding clinical and genomic data from 6909 patients spanning 32 tumor types from the Cancer Genome Atlas (TCGA)^68^ via the Genomic Data Commons (GDC)^69^. We selected one Formalin-Fixed Paraffin-Embedded (FFPE) diagnostic (DX) whole slide image for each patient. If multiple diagnostic slides exist for the same patient, we only used the DX1 image for encoder training. However, we used all slides for the database indexing.

The CPTAC^47^ dataset comprises 7,263 H&E whole-slide images (WSIs) spanning 13 distinct cancer types. These WSIs were sourced from the TCIA (The Cancer Imaging Archive) collection (https://www.cancerimagingarchive.net/browse-collections/). The majority of WSIs were captured at a resolution of 20X magnification. For those originally acquired at 40X magnification, we converted them to 20X to ensure consistency. The standardized 20X WSIs were then used to construct the index database for cancer-type retrieval. Each WSI was annotated with a single image-level label corresponding to one of the 13 cancer types.

We note that resolution may be defined in two ways: as microns per pixel (MPP) or as magnitude, such as 20X, 40X, or 80X. Although MPP is an absolute measure that can be compared across microscopes and databases, most studies use the magnitude definitions, which are not comparable across microscopes. For NCI Lab of Pathology data, either description may be used across all samples because all images were scanned in the same department by the same type of microscope. However, for TCGA and Visium HE images, there is variation and the MPP is not available for many cases. Thus, we used “20X” as an approximation to describe the absolute resolution of the images used to train HERE.

#### 2 Gene mutation

We downloaded the TCGA gene mutation calling results from the GDC^69^. To annotate each mutation’s functional effects, we utilized the OncoKB Annotator^44^ to extract mutation effects from the mutation annotation format (MAF). Subsequently, we categorized gene mutation effects, including “gain-of-function,” “likely-gain-of-function,” “likely-loss-of-function,” “unknown,” or “nan” values, into three categories. For tumor suppressors, we assign categories “Gain_or_unknown”, “Loss”, and “Other” as 0, 1, and 2, respectively. For oncogenes, we assign categories “Loss_or_unknown”, “Gain”, and “Other” as 0, 1, and 2, respectively. As different cancer types have varying gene mutation frequencies, we selected frequently mutated genes for classification tasks, including *CDH1*, *GATA3*, *PIK3CA*, *TP53*, *KRAS*, *ARID1A*, *PTEN*, *BRAF*, *APC*, *ATRX*, and *IDH1*.

For the external evaluation of gene mutation prediction using the CPTAC data, labels were derived from whole-exome sequencing (WXS, Mutation_BCM_v1.zip) datasets from the NCI Proteomics Data Commons (https://proteomic.datacommons.cancer.gov/pdc/cptac-pancancer). These labels were generated following the same processing pipeline used during the training phase with TCGA data.

#### 3 Gene expression

We downloaded the Fragments Per Kilobase of transcript per Million mapped reads (FPKM) data from the GDC^69^. Further, we transformed data to log2(FPKM+1) and subtracted expression values of tumor cases from the mean of normal cases or all cases if normal controls are unavailable. For cases with multiple expression profiles, the median expression values were utilized. These values were further used to compute the mean expression values of the 50 MSigDB Hallmark genesets^58^. Additionally, we derived a cytotoxic T lymphocyte (CTL) value for each case from the median of five genes: *CD8A*, *CD8B*, *GZMA*, *GZMB*, and *PRF1*. Further, we computed four gene signatures related to tumor immune dysfunction and exclusion (TIDE), including cancer-associated fibroblast (CAF), Macrophage M2, myeloid-derived suppressor cell (MDSC), and T-cell dysfunction, following the procedure in the TIDE study^46^. Together, each case had 55 gene expression values as prediction tasks.

For the external evaluation of gene set regression tasks using the CPTAC data, RNA sequencing (RNA-Seq, RNA_BCM_v1.zip) datasets were obtained from the NCI Proteomics Data Commons (https://proteomic.datacommons.cancer.gov/pdc/cptac-pancancer).

For CPTAC, 809 patient cases with both WXS and RNA-Seq data were included in the final performance evaluation. It is important to note that omic data was collected at the patient level, with individual patients often associated with multiple WSIs. Consequently, the same labels were assigned to all WSIs corresponding to a single patient.

### Prediction model design

This section corresponds to the “Stage 1 - Molecular prediction” in Extended Data Fig. 1 and Extended Data Fig. 3a. For an input FFPE diagnostic image, we followed the segmentation pipeline described in CLAM^31^ to extract tissue regions from whole-slide images (WSIs). Specifically, each WSI was first converted from the RGB to the HSV color space. Next, we applied median blurring to the saturation channel to smooth the edges. Based on thresholding segmentation of the saturation channel, a binary mask was generated to delineate the tissue regions. To further refine the mask, we performed an additional morphological closing operation to fill small gaps and holes.

Following the tissue segmentation, we extract non-overlapping image patches with a fixed size (256 x 256 pixels). Each image patch corresponds to an “instance” in multi-instance learning^31^. We used a pre-trained image encoder with frozen network weights to extract features for each patch. Each image was represented by a 𝑁×𝐷 tensor, with the number of patches 𝑁 and the feature dimension 𝐷. This patch-level feature representation is input to a multi-task attention-based multiple-instance learning network with three components: the feature projection module, the gated attention module, and the prediction projection module^31^ (Extended Data Fig. 3a).

The feature projection module stacks multiple fully connected layers to map the input high-dimensional features into a lower-dimensional space. We simplify this module with a single fully connected layer, resulting in a feature dimension 256. For instance, if the backbone feature encoder generates a feature vector with a dimension of 512, the projection layer will have 512 × 256 weights and produce a new feature vector with a dimension of 256. These features are then passed through the gated attention module for further processing.

Given the patch-level embedding tensor *H*ϵ*R*^*N*×256^, the gated attention module assigns attention scores to each patch^31^, generating a new weighted feature tensor *R*^256^. Patches with higher attention scores will have a greater impact on specific tasks than those with lower attention scores. The attention map comprises three fully connected layers with weights *W_a_* ϵ *R*^1×256^, *V_a_* ϵ *R*^256×256^, *U_a_* ϵ *R*^256×256^. Then, for a patch vector *h_n_* ϵ *R*^1×256^, i.e., the n-th row of 𝐻, the attention score for each patch 𝑠_*n*_ ∈𝑅 is computed as follows:

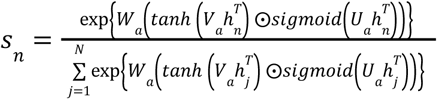

The symbol ⊙ represents the element-wise product between two vectors, which produce a new vector with the same dimension. The attention map operation aggregates the patch-level vectors into one patient-level representation *h_patient_* ϵ *R*^256^ as:

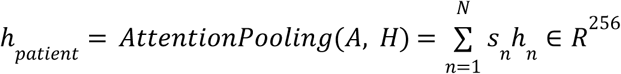

Finally, the prediction layer uses the patient-level feature vector to predict the outputs of multiple tasks. All prediction layers shared the same patient-level features and attention map.

The gene mutation classification tasks cover eleven representative genes, each with three classification labels. Consequently, we have ten prediction layers, each with a weight size of *R*^256×3^.

### Model training

This section corresponds to the “Stage 1 - Molecular prediction” in Extended Data Fig. 1 and Extended Data Fig. 3a. Given the prediction class imbalance across different labels for various tasks, we adopt a weighted cross-entropy loss function for training. The class weights were computed offline for each task based on the class frequency in the training dataset. For the gene expression prediction tasks, we have 55 prediction layers corresponding to the average expression of 50 Hallmark gene sets and five TIDE scores. Each prediction layer has network weights 𝑅^256×1^ for predicting gene expression values. For each regression task, we adopt the mean squared error (MSE) loss function. The total training loss is simply the sum of all these 66 tasks (11 gene mutation classification tasks and 55 gene set regression tasks), defined as follows:

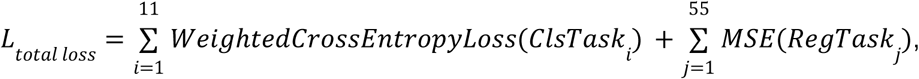

For each classification task, as class distributions were often imbalanced, we employed a weighted cross-entropy loss. The weight for each class c was computed as w_c = 1/p_c, where p_c is the fraction of instances belonging to class c in the dataset. This yields relative weights proportional to the inverse class frequencies, such that less frequent classes receive larger weights and thus exert greater influence on the loss.

We partitioned the TCGA data (6909 cases) into three subsets: train (80%), validation (10%), and test (10%). The splits were constructed at the patient level, ensuring that all slides corresponding to a particular patient were exclusively present in the train, validation, or test set. Meanwhile, splits were repeated until all train/validation/test partitions had positive labels in all gene mutation classes. All image patches (256 x 256) were then resized and z-normalized (by mean and standard deviation) according to different models’ settings.

The training set was utilized to train the deep multi-task attention multiple instance learning network (introduced above), while the validation set aided in selecting optimal hyperparameters, including learning rate, weight decay, and batch size. The test subset was employed to evaluate the model’s performance. Once optimal parameters were determined, we merged the train and validation subsets to train another deep multi-task attention model and selected the best epoch on the test subset. We repeated the train/validation/test split experiments in five independent runs and selected the best model according to the test performance. This model serves as the ultimate encoder for feature extraction in subsequent retrieval tasks.

Via the grid search technique, the optimization hyperparameters were determined as Adam optimization^70^ with a learning rate of 1e-4, b1 coefficient of 0.9, b2 coefficient of 0.999, and L2 weight decay of 1e-5 for 100 epochs. Due to variations in the number of patches per whole-slide image from different patient samples across the whole dataset, the batch size was set to 2 (2 GPUs with mini-batch 1 per GPU). To improve the efficiency and stability of model training, we adopted the gradient aggregation technique, where the gradients across eight sub-batches of data were combined and averaged to imitate the large-batch training.

### Comparison of pre-trained encoders on prediction performance

This section corresponds to the “Stage 1 - Molecular prediction” in Extended Data Fig. 1 and Extended Data Fig. 3c-e.

The HERE prediction framework may utilize different backbone networks for feature extraction. We incorporated four top pre-trained backbones (as evaluated in Extended Data Fig. 2a) as feature extractors with pre-trained weights. The extracted features from these frozen pre-trained models were given as inputs to the multi-task attention-based multiple instance learning framework. In the evaluation, we report the mean and standard deviation of the area under the receiver operating characteristic curves (AUC) for mutation classification tasks and the Spearman correlations for gene expression regression tasks in test datasets (Extended Data Fig. 3c, d). The overall score is the sum of all Spearman correlations for 55 regression tasks and AUC scores for 11 classification tasks.

### Feature vector indexing

This section corresponds to the “Stage 2 - Database indexing” in Extended Data Fig. 1 and Fig. 2a. We utilized feature indexing methods in the Facebook AI Similarity Search (FAISS) library^50^, which implemented various algorithms to reduce memory space and accelerate similarity comparisons for massive amounts of feature vectors. Algorithms utilized in this study include Product Quantization (PQ)^53^, Hierarchical Navigable Small World (HNSW)^51^, Inverted File Index (IVF)^52^, Iterative Quantization (ITQ)^54^, Locality Sensitive Hashing (LSH)^55^. In our evaluation, we studied two representative combinations: ITQ-LSH and HNSW-IVFPQ. Finally, we selected the HNSW-IVFPQ as our database indexing method (Fig. 2). The description below was adapted from the FAISS library documentation^50^:

**HNSW-IVFPQ** is a hybrid indexing approach that combines two techniques: Inverted File (IVF)^52^ with Product Quantization (PQ)^53^ and Hierarchical Navigable Small World graphs (HNSW)^51^. Below, we provide a brief introduction to its implementation. Further algorithmic details are available in each reference cited above.

PQ is a data compression technique that reduces the dimensionality of a feature embedding to speed up the search procedure. It divides a large, high-dimensional vector of a given size into equally sized sub-vectors. Each sub-vector is assigned a “reproduction value” that maps to the nearest cluster centroid of points for that sub-vector. The reproduction values are then assigned to a codebook using unique identifiers, which can be used to reconstruct the original vector.

IVF_PQ is a composite indexing method for approximate nearest neighbor search. It involves dividing the dataset into clusters and quantizing each cluster using product quantization (PQ). Inverted files (IVF) are then built for each quantized cluster, storing references to the original data points. The original data is divided into Voronoi cells, which are partitions that consist of all the points in the space that are within a threshold distance of the given region’s seed point. These seed points are initialized by running K-means over the stored vectors. K-means centroids are turned into the seed points that each define a region. These regions are then used to create an inverted index that links each centroid with a list of vectors in the space, restricting a search to just a subset of vectors.

HNSW aims to locate the nearest Voronoi partition, thereby reducing the search scope. This process entails constructing an HNSW index on all Voronoi centroids. When presented with a query, HNSW is utilized to identify the nearest Voronoi centroid, avoiding a brute-force search method. Subsequently, the query vector undergoes quantization within the relevant Voronoi partition, and distances are computed using PQ codes. HNSW builds a hierarchical multi-layered graph structure where each node represents a data point, and connections are established to other points in the graph based on their similarity. The graph structure allows for fast and efficient exploration of the search space to find approximate nearest neighbors.

Combining HNSW with IVFPQ in FAISS leverages the strengths of both methods: the efficient exploration of the search space provided by HNSW, and the compact representation and fast search capabilities offered by IVFPQ. This hybrid approach can improve performance in large-scale nearest neighbor search tasks, making it suitable for applications such as image retrieval, recommendation systems, and natural language processing.

### Distance metric selection for database indexing

This section corresponds to the “Stage 2 - Database indexing” in Extended Data Fig. 1 and Fig. 2a. The FAISS package that we utilized for database indexing only provides Euclidean distance. However, in our implementation, the Euclidean distance and cosine similarities are monotonically related. We have scaled image encoding vectors to a unit length while preserving their directions, a common pre-processing step in machine learning analysis related to distance calculation and clustering^71^. Then, the cosine similarity is related to Euclidean distance when norms of x and y are 1:

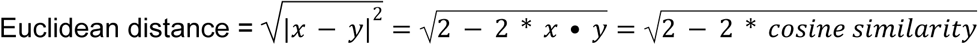

Thus, the top-K results from Euclidean distance and cosine similarity are identical.

### Performance evaluation of vector indexing on image retrievals

This section corresponds to the “Stage 2 - Database indexing” in Extended Data Fig. 1 and Fig. 2b, c.

First, the HERE encoder trained in the deep multi-task attention-based model encoded all image patches into vectors. Then, ITQ-LSH and HNSW-IVFPQ indexing procedures with different parameters compacted vectors into compact codes organized in indexing structures. For ITQ-LSH, we select the code lengths as 32 or 64, leading to 4 or 8 bytes of binary codes, respectively. For HNSW-IVFPQ, there are two critical parameters: the number of nearest neighbors *M* that each vertex will connect to in an HNSW graph, the number of inverted lists *nlist* (i.e., the number of centroids in K-means clustering for data space partition), and the number of sub-quantizers *m* in PQ, which determines the code size and the memory requirement of the final database index. We selected the *M* = *m* in {16, 32}, and *nlist* in {128, 256}.

We evaluate the performance of different indexing approaches and parameter settings on image patch retrieval following the same procedure in the Method section: “*Average precision for comparing image encoders*”. Briefly, for each dataset, the leave-one-out retrieval was conducted. Given one query image, we retrieved the top five results from the indexing database. Then, we report the mean average precision for each class on each dataset. The baseline approach for the optimal accuracy is the brute-force retrieval using the cosine similarity (equivalent to Euclidean distances in our implementation, Methods) among original image vectors.

### Statistical significance computation for top hit patches

This section corresponds to Fig. 1b - Retrieval - Image-to-Image. For each database cohort, such as TCGA or NCI Lab of Pathology data, we randomly selected 10000 image patches to estimate the background distance distribution for a query image. For each top patch hit, we computed the Euclidean distance between the vector encoded from the input image and all randomly selected patches as a group of random distances. The z-score is computed as [distance(input query, top patch) - mean(random distances)] / standard deviation(random distances). The p-value is computed as fractions of random distances <= distance(input query, top patch).

### HERE101 benchmark dataset

This section corresponds to the “Stage 3 - Pathologist scoring” in Extended Data Fig. 1, Fig. 3, and Fig. 4. For the evaluation of retrieval results from web portals, we designed a benchmark HERE101 that comprises 101 diagnostic H&E images from different human organs or tumors (Fig. 3), such as kidney, liver, stomach, lung, breast, colon, ovary, cervix, nerve, lymph node, muscle, soft tissue, and tumors from these sites. Among these images, we included six feature aspects: tumor/normal contents, cell types, tissue sites, tissue structures, cell shapes, and cell cytoplasm. Cell types include epithelial, lymphocyte, mesenchymal, and germ cells. Tissue sites include the lung, liver, ovary, stomach, breast, lymph node, soft tissue, testis, kidney, colon, and others (less frequent sites). Tissue structures include cord, nest, cribriform, sheets, microcystic, swirl, glandular, trabecular, bundle, stratified, and papillary. Cell shapes include polygonal, columnar, cuboidal, round, spindle, and oval. Cytoplasm types include dichroic, basophilic, eosinophilic, clear, and scant.

### Blinded pathologist evaluation of retrieval engines

This section corresponds to the “Stage 3 - Pathologist scoring” in Extended Data Fig. 1, Fig. 3, and Fig. 4. We compared the performance of five competing methods, including PLIP, Yottixel, RetCCL, SISH, and HERE, based on clinical pathologists’ blinded evaluations. The query input to each framework came from the HERE101 benchmark. The output from each framework is the top five whole-slide images (WSI) with the framework name removed and reindexed as methods numbered from 1 to 5.

As the outputs from the five frameworks all have different configurations, we unified the results as follows. For Yottixel, RetCCL, and SISH, we selected the output of the top five WSIs with the most representative image patch per WSI. As HERE outputs multiple image patch hits per WSI returned, we only selected one patch per WSI with the minimal Euclidean distance of encoding vectors from the input query. After these steps, the output configurations of Yottixel, RetCCL, SISH, and HERE are indistinguishable. The output of PLIP is Twitter pages embedded with Twitter-style images, which look very different from WSI. Thus, it is challenging to unify the PLIP’s result configuration with other approaches, and we just removed the PLIP name from the result.

Each pathologist independently assessed the retrieval results of five competing methods (PLIP, Yottixel, RetCCL, SISH, and HERE), yielding three independent ratings per retrieval result, per case, for each method. During the scoring procedure, the pathologist visually examined the top five whole-slide images and scored each whole slide on a five-point whole-number scale, based on the similarity to the input image: 1 (dissimilar), 2 (slightly similar), 3 (similar), 4 (highly similar), 5 (clinically indistinguishable). The final ranking is based on the median score across three pathologists.

### Implementation of previous CBIR frameworks from source code

This section corresponds to the “Stage 3 - Performance evaluation” in Extended Data Fig. 1, Fig. 4, and Fig. 5. We evaluated the retrieval performance of HERE in comparison with four state-of-the-art methods, including Yottixel^19^, SISH^21^, RetCCL^23^, and PLIP^27^, on the HERE101 dataset. Only PLIP provides a public web portal that can be immediately accessed. The other three methods, Yottixel, SISH, and RetCCL, only provide source code, requiring us to build the database locally. The images in the search database were from the NCI Lab of Pathology, which includes 11,232 whole-slide images (WSIs). Each WSI in the dataset is preprocessed by converting it to a 20X image resolution.

The HERE101 benchmark contains 101 high-resolution images, each with a size of 2040 × 1536 pixels at 40X resolution in width and height, corresponding to 1020 × 768 at 20X resolution, as query images. For Yottixel, the patch size in each mosaic of the query image is 1000 × 1000, while for SISH, it is 1024 × 1024. To prepare the input images from HERE101, we first crop the center of each image to 1536 × 1536 and then resize them to 1024 × 1024 to minimize resolution loss. For HERE, we use images of 1020 × 768 at 20X resolution as queries and extract non-overlapping 256 × 256 patches for feature extraction and merging by attention maps.

To evaluate Yottixel, we utilized the source code provided by Kimia Lab with default parameters (https://github.com/KimiaLabMayo/yottixel). First, a mosaic was extracted for each WSI in the search database (i.e., the NCI Lab of Pathology dataset). Each mosaic consists of 5% image patches of all possible patches per whole slide. For every patch, the corresponding Bunch of Barcodes (BoB) feature was extracted. During the query stage, each HERE101 image cropped to 1024 × 1024, was treated as a mosaic containing a single patch. For each query image, we extracted its BoB feature and computed the Hamming distances between the query’s BoB feature and those in the search database. For each WSI in the database, the patch with the minimum Hamming distance was identified. The top-5 WSIs with the smallest Hamming distances were then selected as the retrieval results.

For the evaluation of SISH, we used the official source code at https://github.com/mahmoodlab/SISH. A mosaic was extracted for each WSI in the search database (i.e., the NCI Lab of Pathology dataset) following a similar procedure as Yottixel. Artifact removal was applied to eliminate unwanted elements. Each mosaic consists of 5% image patches of all possible patches per whole slide. Based on the cleaned mosaics, we constructed the SISH index database using the VEB tree. During the query stage, each image in the HERE101 dataset, cropped to 1024 × 1024, was treated as a mosaic containing a single patch. For each query image, we extracted both the VQVAE feature and the binarized DenseNet feature following the same pipeline during the index construction. These query features were then fed into the SISH query pipeline, which returned the top-5 retrieval slides. The returned result for each slide reports one top mosaic patch. Note that, for certain query images, SISH did not return any results when using the default parameters. To address this, we adjusted the Hamming distance threshold to ensure that the retrieval results included at least five WSIs.

For the evaluation of RetCCL, we used the official source code at https://github.com/Xiyue-Wang/RetCCL. Note that only the feature extraction code was provided; for other components, we referred to the source code from another study^72^ (https://github.com/jacobluber/PathologySearchComparison/blob/main/retccl). First, non-overlapping image patches were extracted from each WSI in the search database. RetCCL features were then computed for each patch, and K-Means clustering, based on both RetCCL features and spatial coordinates, was applied to generate mosaics, following the procedure from a previous study^72^, which extracted 20% patches per feature cluster. During the query stage, each image in the HERE101 dataset, cropped to 1024 × 1024, was treated as a mosaic with a single patch. For each query image, the RetCCL feature was extracted, and patch-level retrieval from the search database was performed using linear search with cosine similarities. The top-5 WSIs based on the patch with the smallest distances were returned as the retrieval results.

For the evaluation of PLIP, we used the default database on its official web portal. PLIP encodes each database image as a feature vector of 512 dimensions and performs a linear search for each query image input. This encoding strategy will lead to significant resolution loss for whole-slide images. Theoretically, we can revise the PLIP framework by encoding all individual patches segmented from our whole-slide images (3.2E8 in total from TCGA, CPTAC, and NCI cohorts). However, encoding all patches resulted in an “out of memory” error due to the substantial random-access memory requirement of 5.25 terabytes (3.2E8*512*32 bytes / 1E12 = 5.24288 TB). A similar error occurred when we tried only to encode patches from the NCI Lab of Pathology cohort, which requires two terabytes of random-access memory (122104273*512*32 bytes/1E12 = 2TB).

### Improve the HERE scalability through mosaic sampling

This section corresponds to Fig. 5. The mosaic procedure proposed in Yottixel is an effective approach for reducing the number of processing patches (i.e., the search space) by selecting only the most representative patches from the whole slide image (WSI) prior to database indexing. In contrast, the proposed HERE framework encodes all available tissue patches to retain as much information as possible for the search process. Although the HNSW-IVF-PQ algorithm used in HERE significantly reduces the search space and time through a hierarchical navigable graph and efficient memory optimization via product quantization, further improvements may be achieved by integrating a mosaic-based approach similar to that of Yottixel. To explore this possibility, we conducted an additional experiment incorporating the mosaic procedure into the HERE framework, referred to as HERE(mosaic). In this variant, only the top 20% of representative patches—selected through the mosaic method—are indexed within the HERE system. The mosaic generation process mirrors that of Yottixel, with the selection of the top 20% of patches per WSI based on representativeness, using a patch size of 256 × 256. We evaluated this integrated approach on the CPTAC cohort for both cancer type retrieval and mutation retrieval tasks.

### Development of the public web portal

We first collected a large cohort of H&E image data. The collected H&E data archive contains 30,435 H&E whole slide images from the NCI Lab of Pathology, TCGA^68^, CPTAC^47^, and spatial transcriptomic datasets (Supplementary Table 1).

We first conduct the image segmentation for each whole slide image and extract the image patches within tissue regions (example in Extended Data Fig. 3b). The number of image patches is more than 3.2E8. Then, the HERE encoder converts all image patches as vectors. The HNSW-IVFPQ framework further organized all vectors from the NCI Lab of Pathology, TCGA, and CPTAC, into three database indexes. Finally, all database indexes together take about 12.1 gigabytes, which can be loaded into modern computer memory.

We used the Flask (v3.0.0) framework based on Python (3.9) language for the backend development, HTML and JavaScript for the frontend development, the Apache HTTP server (v2.4.6) with WSGI module (v5.0.0), MySQL database (v8.4.0) for web service.

### Computational resource specifications

We listed the resources allocated for key computational parts. The retrieval is done online in real-time, thus requiring high processing speed. All of the other calculations are pre-computed offline and thus are less time-sensitive. As input data may vary, we provide only what we consider a typical example when reporting the running time for each part.

Retrieval (The core function of HERE):

● CPU: 1 Intel Xeon Platinum 8259CL from Amazon Web Service (AWS).
● GPU: 0
● MEM: 64GB. The database indices from three large H&E cohorts (NCI, TCGA, CPTAC) take 12.1 GB. Thus, the actual memory used is less than the upper limit.
● DISK: shared disk storage on Amazon Web Services (AWS) with a five-terabyte quota. The largest disk storage consumption is the storage of whole slide images for result visualizations. To fit our AWS quota, we downsampled the NCI, TCGA, and CPTAC images to 4.1 terabytes. However, we finished the encoding and database indexing steps in the NIH high-performance computing cluster, thus did not downsample the image and use 20X resolution because we store original images on the NIH cluster.
● Running time: See Fig. 2d for TCGA and NCI cohorts.

Database indexing:

● CPU: 4 AMD EPYC 9454 from the NIH BioWulf high-performance computing cluster.
● GPU: 0
● MEM: 100GB
● DISK: shared disk storage on the NIH Biowulf cluster with 25TB quota
● Running time: 28.14 minutes for indexing the NCI Lab of Pathology data with 122,104,273 feature vectors in 256 dimensions.

Training of encoder:

● CPU: 4
● GPU: 2 Nvidia V100x
● MEM: 64 GB
● DISK: shared disk storage on the NIH Biowulf high-performance cluster with a 25TB quota
● Running time: 27 hours 55 minutes for training 100 epochs of CONCH backbones on TCGA data.

Inference of prediction labels on whole-slide images:

● CPU: 4
● GPU: 1 Nvidia V100x
● MEM: 32GB
● DISK: shared disk storage on NIH Biowulf with 25TB quota
● Running time: average time: 33.19 seconds, standard deviation 22.76 seconds on average for whole slide images from CPTAC, with an average size of 1.74 E9 pixels in 20x.

### Query image features from gene transcriptomics input

This section corresponds to the “Stage 4 - Transcriptomics - image feature association” in Extended Data Fig. 1 and Fig. 6a-d. For each spatial transcriptomics (ST) data paired with an H&E image, we first extracted patches centered around ST detection spots with the patch size as 2^ceil(log(64/micron-per-pixel)/log(2)). Then, we utilized the HERE patch encoder to convert image patches into vectors in 256 dimensions. We applied clustering to organize the image patches centered around ST detection spots into 8 clusters based on the cosine similarities (equivalent to Euclidean distances in our implementation, Methods) between encoded vectors and the k-means algorithm. For each gene in each cluster, we performed the two-sided Wilcoxon rank-sum test and computed Cohen’s d values between ST spots in the cluster and ST spots outside the cluster. Once a user queries a gene, HERE will scan all ST datasets’ clusters and report image clusters where the gene expression is high with thresholds Cohen’s d > 1 and false-discovery rate (FDR, computed as the two-sided Wilcoxon rank-sum test p-value with the Benjamini-Hochberg correction) < 0.05.

### Mice

This section corresponds to the “Stage 4 - In vivo study” in Extended Data Fig. 1 and Fig. 6e, f. All animal experiments were approved by the NCI Animal Ethics Committee of NIH and performed strictly according to the animal protocol CDSL-001. C57BL/6 mice were purchased from the Charles River Laboratories (NCI strains). Female mice at 6–8 weeks of age were used for subcutaneous tumor incubation. The maximal tumor size permitted by the ethics committee is 2 cm in the longer diameter, or the tumor volume reaches 2000 mm^3^, and no tumor burden exceeds that limit in our experiment. Mice were euthanized by carbon dioxide inhalation when the tumor size reached the endpoint. Mice were housed in 12-hour light-dark cycles with ambient temperatures of 20–25 °C and 40–60% humidity.

### Cell lines and cultures

This section corresponds to the “Stage 4 - In vivo study” in Extended Data Fig. 1 and Fig. 6e, f. The mouse melanoma cell line B16F10 was purchased from the American Type Culture Collection (CRL-6475). B16-mhgp100 was provided as a gift by R. J. Kishton from the Surgery Branch at NCI, NIH. B16 (H-2b), the source of B16-mhgp100, is a gp100+ spontaneous murine melanoma obtained from the NCI tumor repository. B16F10 and B16-mhgp100 cells were routinely cultured in RPMI 1640 medium (Gibco) supplemented with 10% FBS (Gibco BRL) and 100 IU ml−1 penicillin/streptomycin (P/S) and 20 mM HEPES. The 293FT cells (Thermo Fisher, R70007) were cultured in complete medium, which is high-glucose DMEM (Gibco) supplemented with 10% FBS (Gibco BRL), 100 IU ml−1 P/S, 1 mM sodium pyruvate (Gibco), 0.1 mM MEM NEAA (Gibco) and 0.5 mg ml−1 geneticin (Gibco). All cells were incubated in a humidified incubator at 37 °C with a 5% CO2 supply.

### Cell Proliferation Assay

This section corresponds to the “Stage 4 - In vivo study” in Extended Data Fig. 1 and Extended Data Fig. 9e. B16F10 cells were seeded at 1,000 cells/well in a 96-well plate and incubated in a 37 °C humidified CO2 incubator. After overnight incubation, the number of cells was determined by the Cell Proliferation Kit II (XTT, Roche) every day, as per the manufacturer’s protocol. The absorbance was measured at 460 nm with a reference wavelength of 620 nm using a microplate reader.

### RT–qPCR analysis

This section corresponds to the “Stage 4 - In vivo study” in Extended Data Fig. 1. Total RNA was extracted from tumor cells by using the PureLink™ RNA Mini Kit (Invitrogen), and cDNA was synthesized through reverse transcription using the SuperScript™ III First-Strand Synthesis SuperMix for qRT-PCR (Invitrogen). RT–qPCR was performed using the SYBR™ Green PCR Master Mix (Applied Biosystems) and corresponding primers, and Ct values were detected by StepOne Plus Real-time PCR system. Raw data were processed using SDS 19.1 software, and the relative mRNA expression level was normalized to the internal ribosomal reference gene *Rpl13a*. The primer sequences (5’-3’) are listed as follows:

*Rpl13a* Forward: AGCCTACCAGAAAGTTTGCTTAC

*Rpl13a* Reverse: GCTTCTTCTTCCGATAGTGCATC

*Serping1* Forward: CACCAACCATAAGATCCGCAA

*Serping1* Reverse: GGCACTCAAGTAGACAGCATT

### Lenti-viral transduction for *Serping1* over-expression

This section corresponds to the “Stage 4 - In vivo study” in Extended Data Fig. 1. Full-length cDNA of murine *Serping1* were constructed into the GeneCopoeia EF1a Lv195 plasmid. The plasmid DNA was transformed into One Shot™ Stbl3™ Chemically Competent E. coli (Invitrogen) via heat shock, and was then purified with EndoFree Maxi Plasmid kit (TIANGEN), according to the manufacturer’s instruction. 1×10^6^ per well Lenti-X™ 293T cells (Takara) were seeded onto a 6-well plate. 24 hours after cell seeding, two packaging plasmids psPAX2 (Addgene 12260) and pMD2.G (Addgene 12259), together with the Lv195 vector or Lv195-*Serping1* plasmid, were transfected into the 293T cells using the Lipofectamine 3000 transfection reagent (Invitrogen), according to the manufacturer’s protocol. Culture medium was replaced with Opti-MEM™ with GlutaMAX™ (Gibco) supplemented with 1:500 ViralBoost Reagent (Alstem) 6 hrs after transfection. The lentivirus was harvested by spinning down the viral supernatant at 1,000x g, 4°C for 15 mins to remove the cell debris at 48 and 72 hrs post transfection.

For lentivirus transduction, B16-mhgp100 and B16F10 cells were mixed with lentivirus at the 1:1 dilution with culture medium. 10 μg/mL polybrene (Sigma-Aldrich) was added to the mixture for 24-48 hours before refreshing the medium. Two days after infection, puromycin (Sigma-Aldrich, 2ug/ml) was added for the selection and maintenance of cells with positive *Serping1* expression.

### In-vivo validation of *Serping1*’s effects on tumor growth

This section corresponds to the “Stage 4 - In vivo study” in Extended Data Fig. 1 and Fig. 6e, f. For subcutaneous tumor models, 1×10^5^ B16F10 or B16mhgp100 cells were injected subcutaneously into 6-8-week-old female C57BL/6 mice. Anti-PD-1 antibody (200ug per mouse, equivalent to 10 mg/kg with 20g per mice, BioXCell InVivoMAb, Clone RMP1-14) was given intraperitoneally to the mice twice a week for total 4 doses, starting on day 6 (B16F10) or day 10 (B16-mhgp100) post tumor inoculation. Anti-CTLA4 antibody (200ug per mouse, equivalent to 10 mg/kg with 20g per mice, BioXCell InVivoMAb, Clone 9D9) was given to B16F10-bearing mice twice a week for a total of 4 doses, starting on the next day following the 1st PD-1 treatment. Tumor volume was measured based on the formula: (Length×Width^2^)/2. Mice were euthanized according to endpoints defined as tumor volume ≥ 2000 mm3 or the tumor length (longer diameter) ≥ 2 cm.

### Data availability

For data from the NCI Lab of Pathology, all H&E images and available clinical labels are available on the web portal: https://hereapp.ccr.cancer.gov. All TCGA data are downloaded from the GDC https://gdc.cancer.gov. All CPTAC data are downloaded from the TCIA (https://www.cancerimagingarchive.net/browse-collections/) and https://proteomic.datacommons.cancer.gov/pdc/cptac-pancancer. The sources of spatial transcriptomics data are listed in Supplementary Table 1. The HERE101 benchmark is available for download at https://github.com/data2intelligence/HERE101.

The trained encoder model based on the CONCH backbone is available at https://github.com/data2intelligence/HERE_training/blob/main/assets/snapshot_53_HERE_CONCH.pt

### Code availability

The source code of the HERE web portal is available at https://github.com/data2intelligence/HERE_website.

The source code for training the HERE encoder and prediction model is available at https://github.com/data2intelligence/HERE_training.

### Ethics & Inclusion statement

All research activities conducted in this study followed the ethics and inclusion criteria suggested by Springer Nature.

## Acknowledgments

We thank Dr. Seongyong Park for helpful discussions. The triple-negative breast cancer (TNBC) full-resolution H&E images from the Visium platform were kindly provided by Dr. Rania Bassiouni. The work was supported by the NIH intramural budget, the NCI FLEX award, the Technology Impact Award from the Cancer Research Institute (all awarded to P.J.), and Guangdong Basic and Applied Basic Research Foundation (2023A 1515010109, awarded to Y.Z.). This work utilized the computational resources of the NIH HPC Biowulf cluster (https://hpc.nih.gov).

## Contributions

P.J. and Z.Z. designed the study, performed computational analyses, and wrote the manuscript. Z.Z. established the web server. J.H. designed the HERE101 benchmark. J.H., X.L. and L.S. performed web portal evaluations. L.G. performed wet-laboratory experiments. B.R. collected the spatial transcriptomics data. R.P. and J.L. deployed the HERE server under the government domain. K.A. provided the NCI Lab of Pathology data and wet laboratory space. P.J. and Y.Z. provided funding.

### Extended Data Figures

**Extended Data Fig. 1:**
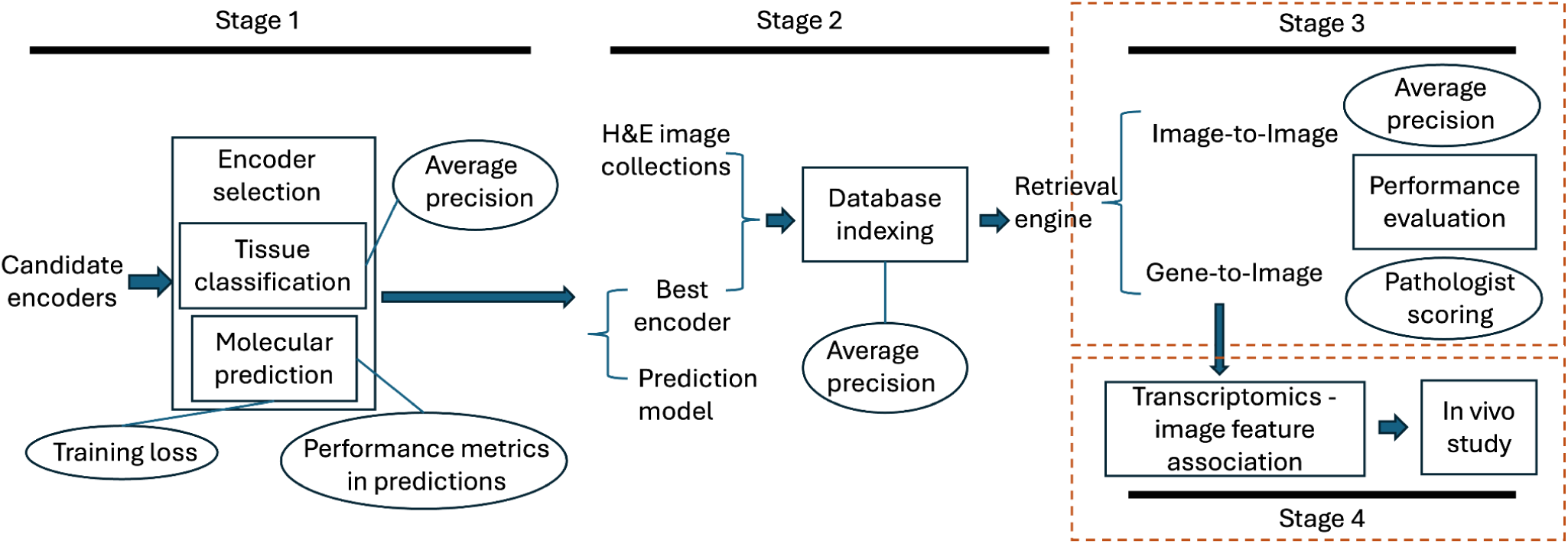
Study flow in four stages. Stages 1 and 2 correspond to the same stages 1 and 2 in Fig. 1a, except that Fig. 1a introduces the structure of prediction and database frameworks. In contrast, this figure introduces the study flow with input and output at each stage. Each box represents an experimental task. Each elliptical node represents an evaluation or training metric utilized in an experiment. Each text without bounding shapes represents the input or output to an experimental task. Arrows indicate the order of experiments performed. The Stage 1 task “Encoder selection” comprises two sub-tasks: Tissue classification & Molecular prediction. For the sub-task molecular prediction, the training of the prediction framework is based on “Training loss” functions (Methods). However, the comparison of encoder backbones are based on “Performance metrics in predictions” (Methods).

**Extended Data Fig. 2:**
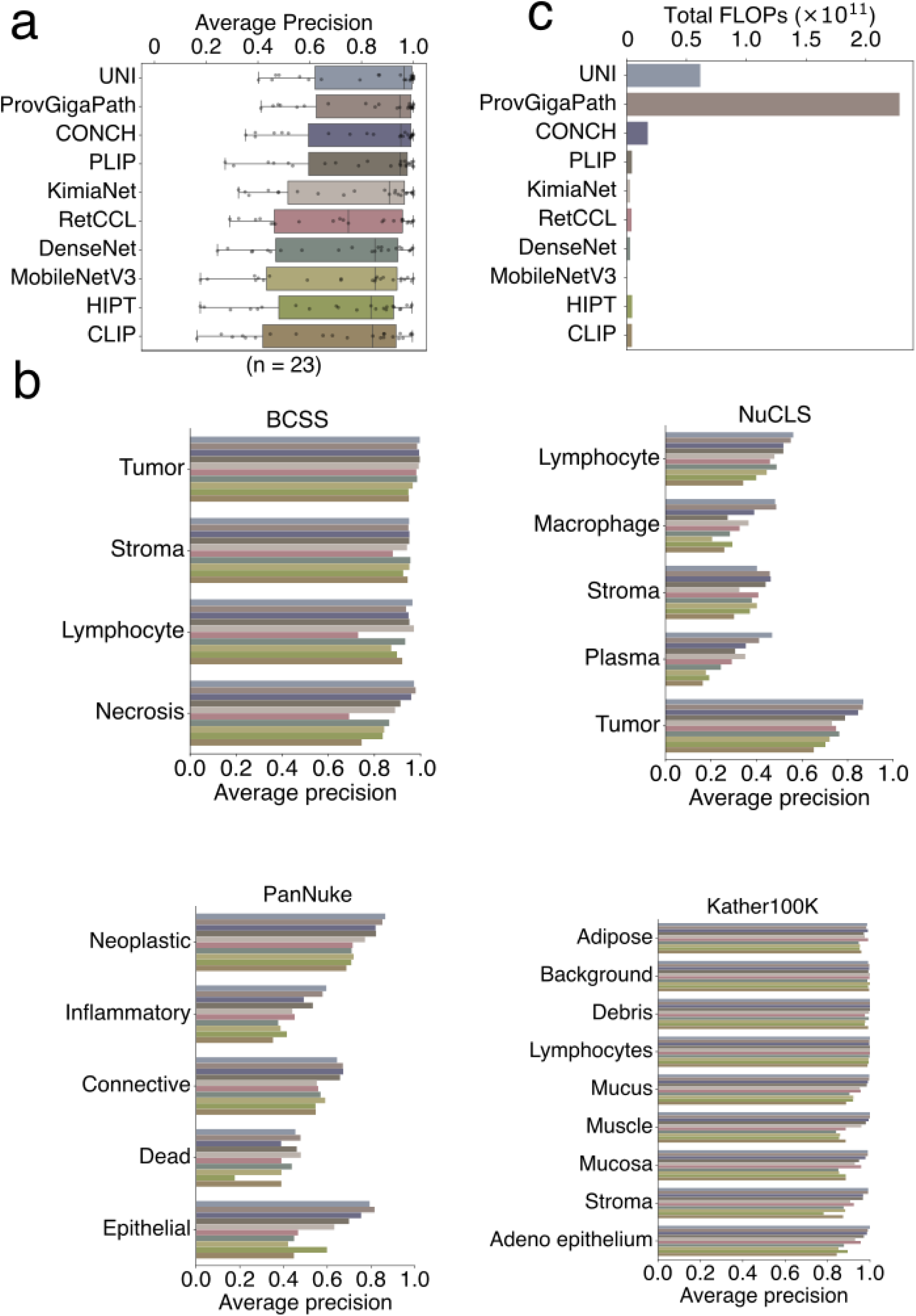
Accuracy comparison among image encoders. **a,** Average precisions of image encoders across all evaluation datasets, across all classes and datasets (details in panel b). The box plots are shown as Fig. 2b. **b**, Accuracy comparisons among seven image encoder approaches in individual datasets. Four H&E annotation datasets, including BCSS, NuCLS, PanNuke, and Kather100K, were used to evaluate the average precision (across all patches) of top-5 results from each image encoder (Methods). **c**, Computational cost quantified as the floating-point operation (FLOP).

**Extended Data Fig. 3:**
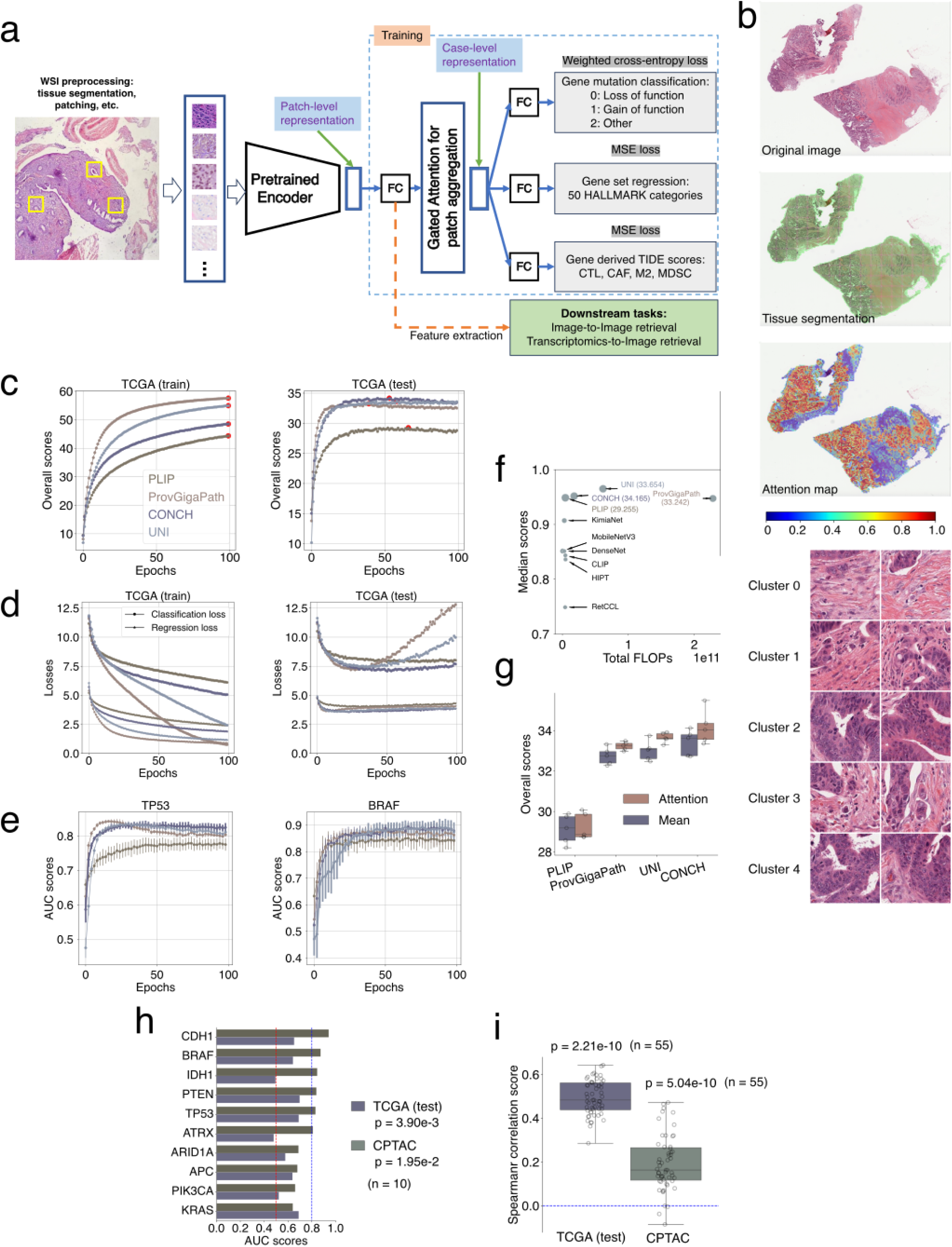
Model training in the HERE framework. **a,** Multi-task multi-instance prediction model. For each whole-slide image (WSI), the training framework applies a coarse color-based segmentation to locate the tissue region and extract non-overlapping image patches. Then, a pre-trained image encoder converts patches to vectors with the same dimension. The HERE framework includes a fully connected network (FC) to convert output vectors from different encoders to the same format of 256 dimensions. Then, a gated attention map aggregates patch vectors from the same slide into one vector, utilized by additional network layers to predict gene omics status (Methods). **b,** Examples of an original image (top) tissue segmentation (middle) and an attention map (bottom). The original image was downloaded directly from TCGA (1ec9435b-6056-4d34-802d-a4d20208f60e). For tissue segmentation and patching, green lines indicate the boundaries between patches (Methods). For the attention map (bottom), colors indicate normalized attention scores, which highlight feature regions to later tasks. High attention scores (red, scale bar) indicate greater significance, with low scores (blue, scale bar) less influential. A few additional image patches with high attention scores were attached to the bottom of this panel (adapted from the case TCGA-G4-6303-01Z-00-DX1). **c,** The training and test curves on the TCGA. Four encoders were evaluated as the pre-trained encoder in panel a. The training procedure followed the train/validation/test split scheme (Methods), but only the train and test curves are shown after finalizing hyperparameters with validation splits. For each encoder, the best epoch (achieving the highest overall score) was labeled by a red dot. The overall score is the sum of all Spearman correlations for 55 regression tasks and AUC scores for 11 mutation classification tasks. The mean scores across five splits are shown. **d,** Loss curves plotted separately for classification tasks and regression tests, as in panel c. **e,** Prediction performance on example outputs, including *TP53* loss-of-function mutation and *BRAF* gain-of-function mutation. Dots and error bars represent the mean and standard deviation across five random data splits. **f**, Encoder metrics summary. The x-axis presents the total floating-point operation (FLOP) as the computational cost for encoders. The y-axis presents the median score from the image-patch classification performance (Extended Data Fig. 2a). For the top four encoders analyzed in panel c, their overall performance scores in the TCGA test data (red dots on the test curve in panel c) were labeled. CONCH was selected as the backbone encoder in HERE due to its high performance metrics and low computational cost. **g**, Ablation experiment replacing the attention-based aggregation module with a simple mean operation. Each dot corresponds to one train/test split. The overall score (defined as panel c) in the test set comes from the best epoch (selected on the TCGA test data) for each method. **h,** Prediction performance of mutation status on TCGA test data and CPTAC independent test cohorts. The area under the ROC curve (AUC) of binary mutation prediction is shown based on the best model trained with the TCGA data. The p-value is from the two-sided Wilcoxon rank-sum test, comparing group values and 0.5, with the Bonferroni correction. **i,** Prediction performance of the average expression of gene sets on held-out TCGA test data and independent CPTAC test cohorts. The Spearman rank correlation between predicted and real values is shown in box plots as Fig. 2b.

**Extended Data Fig. 4:**
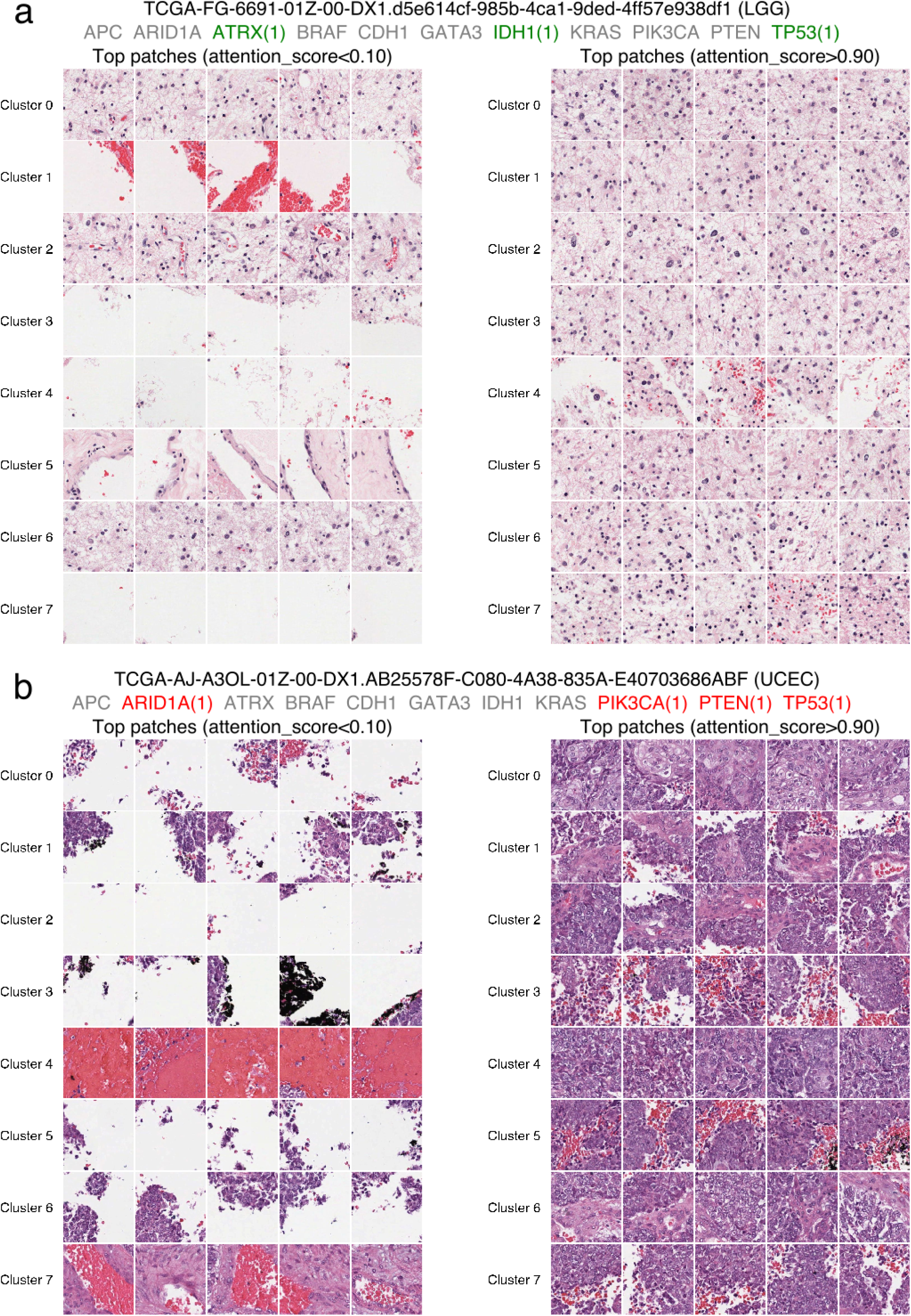
Example attention maps from true and false predictions. In each case, patches with high and low attention scores are clustered based on their encodings and representative clusters are shown. For each whole-slide image, we first normalize the original attention scores into [0,1]. Then, we cluster those patches with high attention scores (>0.9, right) into 8 clusters on the image embeddings. Within each cluster, the top five patches with nearest distances to the center are selected for visualization. The same procedure is conducted for patches with low attention scores (<0.1, left). **a**, Example patches with high and low attention scores from a TCGA test case (Astrocytoma) with three mutations correctly predicted (highlighted in green). **b**, Example patches with high and low attention scores from a TCGA test case (Endometrial carcinoma) with four mutations falsely predicted as negative (highlighted in red).

**Extended Data Fig. 5:**
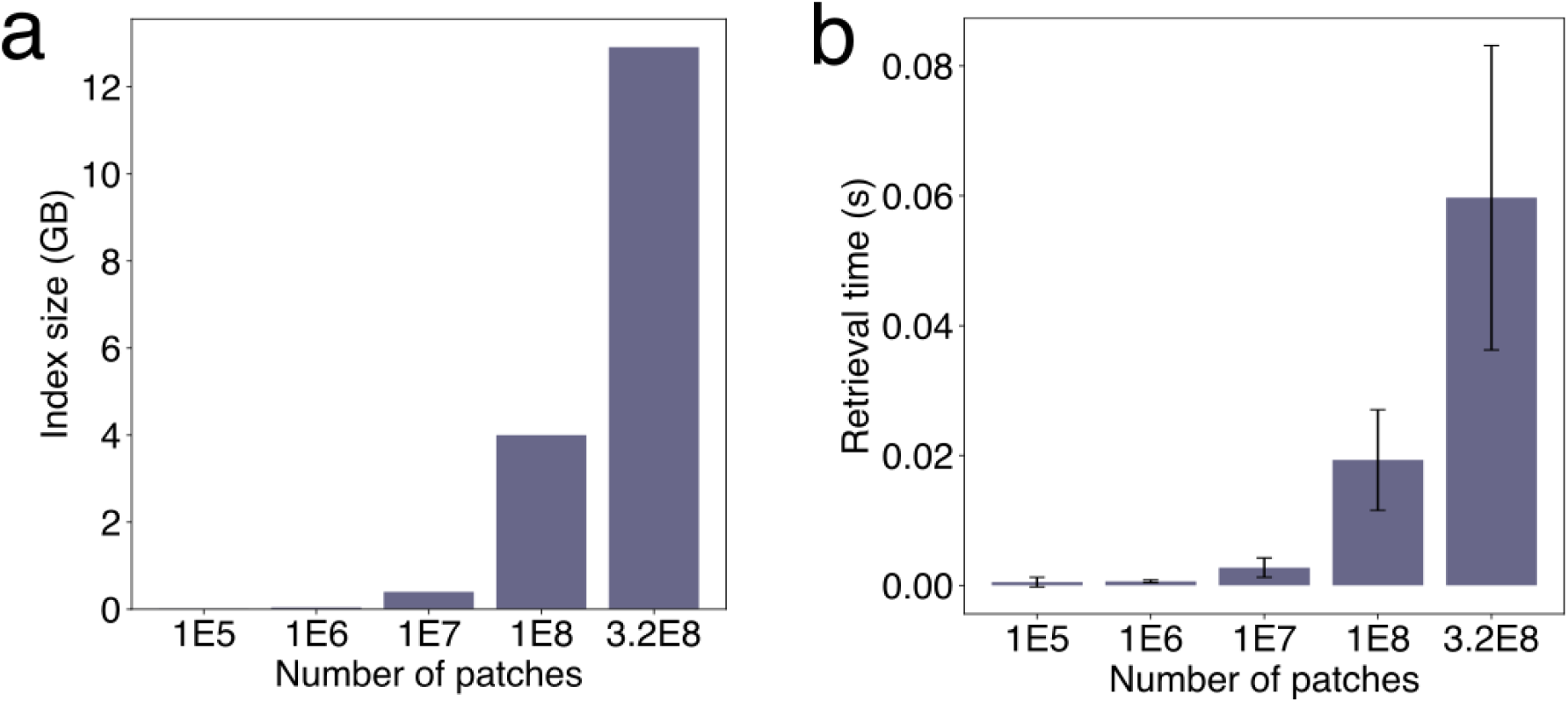
Scalability analysis of the retrieval system. We measured the database index size and retrieval time (excluding other computational parts such as image encoding and network transfer delays) with different numbers of patches indexed in the database. 3.2E8 is the number of patches from all cohorts (TCGA, CPTAC, and NCI). **a**, Index size. **b**, Retrieval time, mean and standard deviations (error bars) across 101 images from the HERE101 benchmark (Fig. 3).

**Extended Data Fig. 6:**
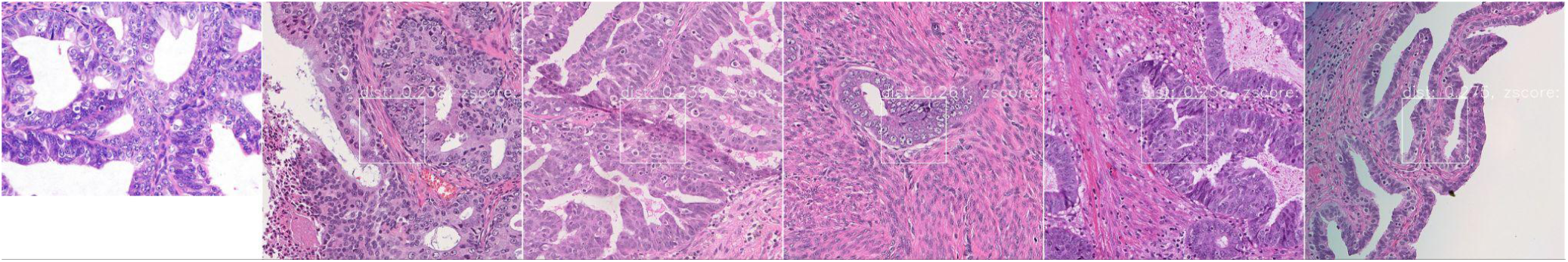
Example of pathologist scoring decisions. The image on the left (**first patch**) is a query image (Case No. 1037428-1.1) from the HERE101 dataset. The five images on the right are the top 5 retrieved patches from five different whole-slide images (WSI) returned by the HERE search engine. Note that HERE returns multiple patches for each WSI. However, in the blinded evaluation, we only selected one top patch per WSI (based on the minimal distance metric) to mask the output difference between HERE and other frameworks. The pathologist #1 evaluated these retrieved patches and assigned them the following scores: 5, 4, 3, 4, and 2. The rationale behind these scores is as follows: ● **Score 5 (second patch):** The glandular cells exhibit cribriform architecture, and the tumor cells are oval-shaped with vesicular nuclei; small nucleoli are also visible. ● **Score 4 (third patch):** Either the architectural or nuclear features are not perfectly matched. **Score 3 (fourth patch):** This patch contains a single glandular structure with a significant presence of non-tumoral components. ● **Score 2 (last patch):** The cells are oval-shaped, bearing some resemblance to the query, but the overall structure is papillary.

**Extended Data Fig. 7:**
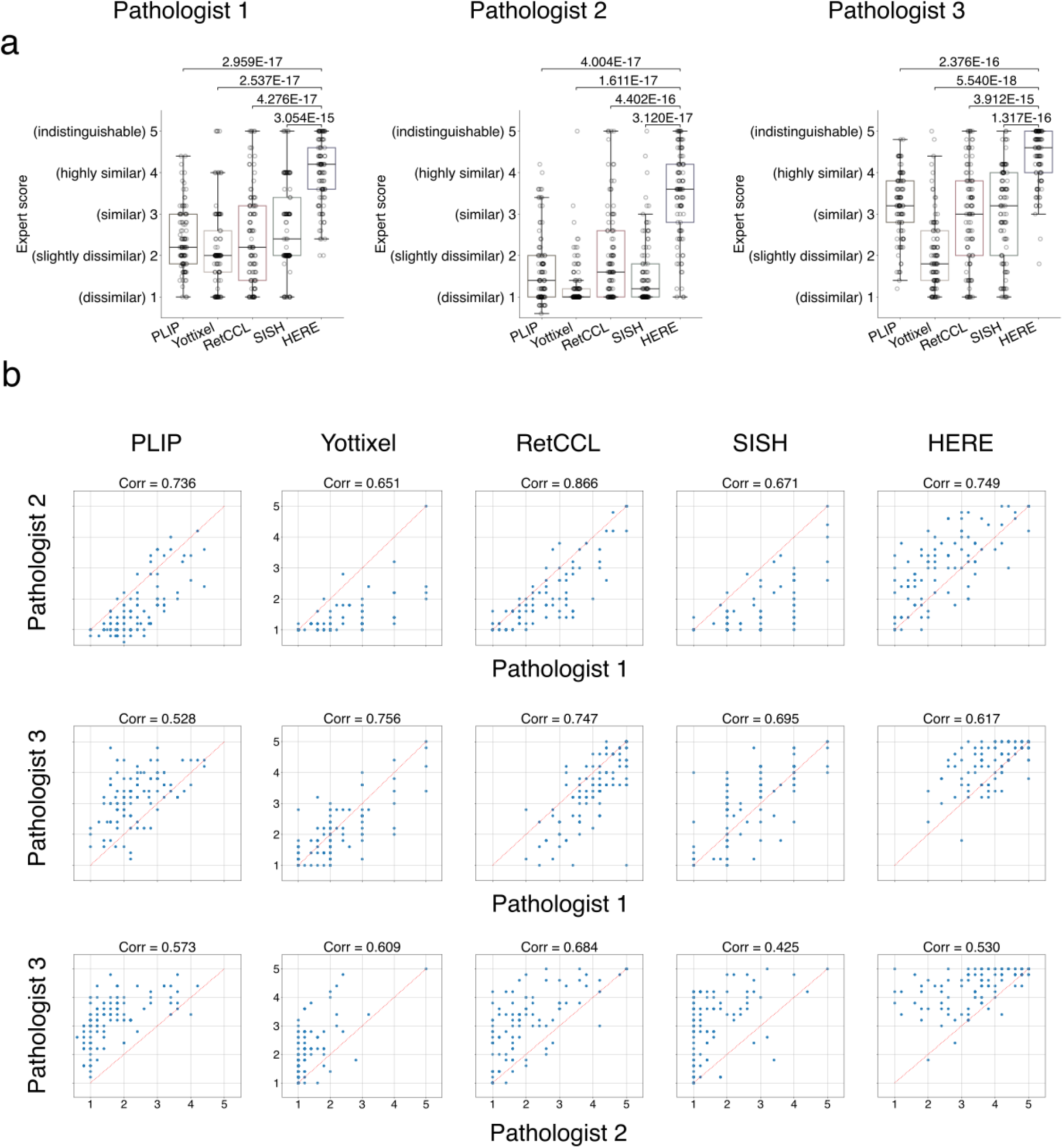
Blinded accuracy assessment on HERE101 benchmark. **a**, Individual plot for each pathologist. Within each box plot, each dot represents the average score among the top 5 whole-slide images for a query (n= 101 per box group). The thick line represents the median value. The bottom and top of the boxes are the 25th and 75th percentiles, respectively (interquartile range). Whiskers encompass 1.5 times the interquartile range. The P-value was calculated using the two-sided Wilcoxon signed-rank test, comparing scores between HERE and other methods paired on each input image. **b**, Correlations of scores from different pathologists. To assess the reliability of the human evaluations, we computed Pearson correlations between three pathologists’ ratings for each method (n = 101 per correction).

**Extended Data Fig. 8:**
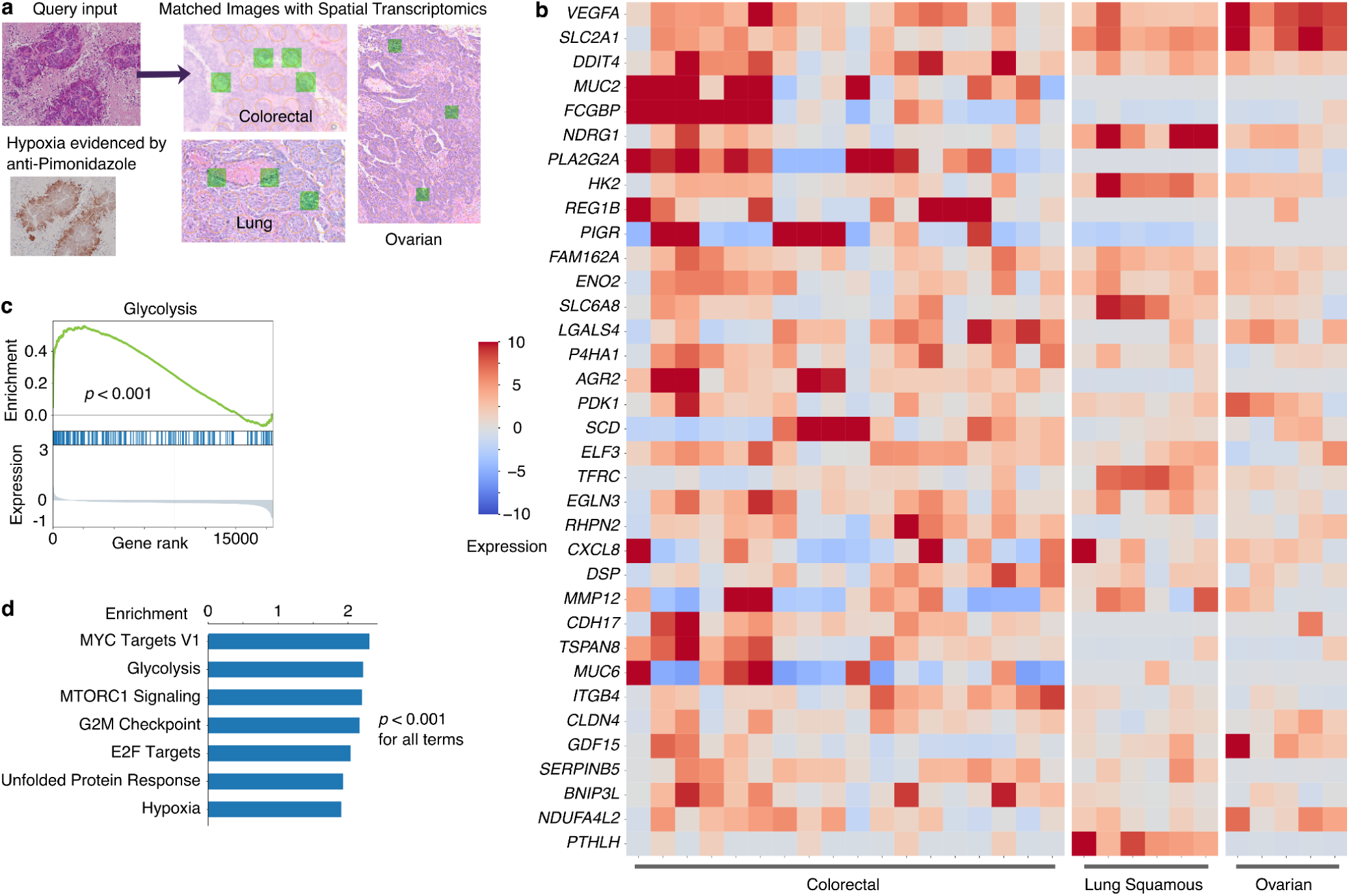
HERE returns gene expression profiles given an input image with spatial transcriptomics. **a**, An example query image from a tumor with hypoxia indicated by anti-Pimonidazole staining^57^. The right side shows examples of matched image patches (green squares) within an H&E slide with spatial transcriptomics (ST) data. **b**, Transcriptomics heatmap of ST detection spots from the top three ST profiles, each representing a distinct cancer type. Expression values are variance-stabilizing transformed values relative to levels in all ST spots and sequencing depth per slide^73^. Only genes with expression values greater than 5 in at least three spots across all match patches are shown, and genes are ranked by mean values across all spots. Within each cancer type, columns (spots) are organized by hierarchical clustering with correlation distances. Only spots with expression values greater than 5 in at least three genes are shown. **c**, Glycolysis gene set enrichment. Along the x-axis, all genes are ranked from high to low by mean expression value (lower y-axis) among all ST detection spots returned by the image query. Members of the glycolysis hallmark gene set^45^ are indicated by horizontal blue lines in the middle of the plot. The top y-axis plots the glycolysis enrichment score at each gene rank. The p-value was computed through the one-sided permutation test with 1000 randomizations. **d**, Gene sets with higher enrichment scores than hypoxia.

**Extended Data Fig. 9:**
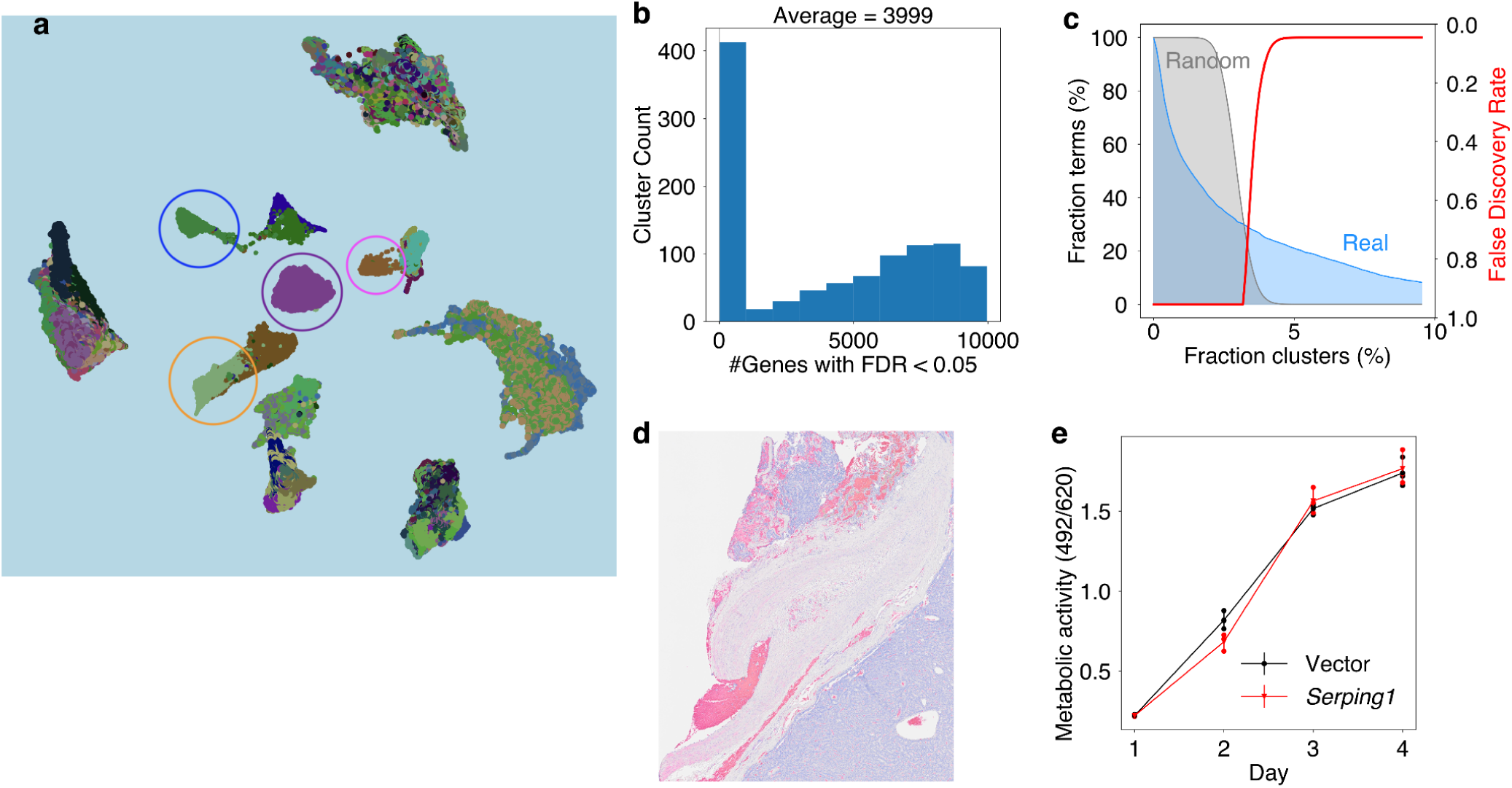
Image features associated with gene expression levels. **a**, Screenshot of the 3D UMAP of patch encoding vectors from H&E images paired with spatial transcriptomics. The full interactive UMAP is available at https://hereapp.ccr.cancer.gov/ST_CONCH_umap3d.html. Some image patch clusters or local regions comprise image patches from mostly one or two ST profiles (circle highlights), typical of batch effects. **b**, Statistical links between gene expression and image feature clusters. For each ST cluster, we counted the number of genes with FDR < 0.05 (Benjamini-Hochberg corrected from the two-sided Wilcoxon rank sum test). The histogram of gene count values across image clusters from all ST profiles is shown. The total number of clusters evaluated was 1,039, and the total number of genes was 11,137. **c**, Gene set enrichment analysis. For Cohen’s d profile for each ST cluster, we performed gene set enrichment analysis. The X-axis presents the fraction of ST clusters above which a GO_BP term is enriched (GSEA q-value < 0.05). The left Y-axis presents the fraction of GO_BP terms enriched above the threshold on the X-axis for real and randomly permuted data. The right Y-axis presents the False Discovery Rate, computed as the (Random GO_BP term fraction) / (Real GO_BP term fraction). **d**, The H&E image from the human lung tumor region shown in Fig. 6a, which has high expression of *C1R*, *C1S*, and *SERPING1*. **e**, In-vitro growth of B16F10 cancer cells in culture, measured by XTT assay. Dots and error bars represent mean and standard deviations (n = 3 cell culture replicates). Metabolic activity is measured as optical density at 492 nm (read) divided by the value at 620 nm (reference)

### Supplementary Tables

**Supplementary Table 1:** Catalog of tumor spatial transcriptomics profiles paired with HE images. Available as file “Supp_Table_1.xlsx”

**Supplementary Table 2:** Cancer functional associations of top enriched pathways.

|  | Interpretation |
| --- | --- |
| Cytoplasmic Translation | Deregulation of protein translation is a common feature of cancer and a target of anticancer therapy<sup>74</sup>. |
| Antigen Processing & Presentation | Involved in anti-tumor immune response by T cells. |
| Humoral Immune Response | Anti-tumor immunoglobulin responses may antagonize cancer growth<sup>75</sup>. |
| Collagen | Collagen deregulation can promote cancer progressions<sup>76</sup>. |
| Adaptive Immune Response | Involved in anti-tumor immune response through the adaptive immune cycle. |
| External Encapsulating Structure | External encapsulating structures, such as fibrotic stromas, are common morphologies in tumors<sup>76</sup>. |
| Leukocyte Mediated Cytotoxicity | Involved in anti-tumor immune response by cytotoxic cells killing tumor cells. |
| Leukocyte Mediated Immunity | Involved in anti-tumor immune response. |
| Cell Killing | Involved in anti-tumor immune response by cytotoxic cells killing tumor cells. |
| Complement Activation | The complement system's role in cancer is controversial and can act as either a negative or positive regulator of tumorigenesis<sup>64</sup>. |
| Immune Effector Process | Involved in anti-tumor immune response. |

**Supplementary Table 3:** Data availability summary for multimodal datasets. The tables list the maps between tumor case ID, image file names, availability of mutations, and transcriptomics, utilized in the encoder training. Available as file “Supp_Table_3.xlsx”

## Notes

### Competing Interest Statement

The authors have declared no competing interest.

## References

1. Zheng, L., Wetzel, A. W., Gilbertson, J. & Becich, M. J. Design and analysis of a content-based pathology image retrieval system. IEEE Trans. Inf. Technol. Biomed. 7, (2003).

2. Qi, X. et al. Content-based histopathology image retrieval using CometCloud. BMC Bioinformatics 15, 287 (2014).

3. Zhang, X., Liu, W., Dundar, M., Badve, S. & Zhang, S. Towards large-scale histopathological image analysis: hashing-based image retrieval. IEEE Trans. Med. Imaging 34, 496–506 (2015).

4. Sridhar, A., Doyle, S. & Madabhushi, A. Content-based image retrieval of digitized histopathology in boosted spectrally embedded spaces. J. Pathol. Inform. 6, 41 (2015).

5. Kwak, J. T., Hewitt, S. M., Kajdacsy-Balla, A. A., Sinha, S. & Bhargava, R. Automated prostate tissue referencing for cancer detection and diagnosis. BMC Bioinformatics 17, 227 (2016).

6. Sparks, R. & Madabhushi, A. Out-of-Sample Extrapolation utilizing Semi-Supervised Manifold Learning (OSE-SSL): Content Based Image Retrieval for Histopathology Images. Sci. Rep. 6, 27306 (2016).

7. Jiang, M., Zhang, S., Huang, J., Yang, L. & Metaxas, D. N. Scalable histopathological image analysis via supervised hashing with multiple features. Med. Image Anal. 34, 3–12 (2016).

8. Shi, X. et al. Supervised graph hashing for histopathology image retrieval and classification. Med. Image Anal. 42, 117–128 (2017).

9. Ma, Y. et al. Breast Histopathological Image Retrieval Based on Latent Dirichlet Allocation. IEEE J Biomed Health Inform 21, 1114–1123 (2017).

10. Zheng, Y. et al. Histopathological Whole Slide Image Analysis Using Context-Based CBIR. IEEE Trans. Med. Imaging 37, 1641–1652 (2018).

11. Akakin, H. C. & Gurcan, M. N. Content-based microscopic image retrieval system for multi-image queries. IEEE Trans. Inf. Technol. Biomed. 16, 758–769 (2012).

12. Zheng, Y., Jiang, B., Shi, J., Zhang, H. & Xie, F.-Y. Encoding Histopathological WSIs Using GNN for Scalable Diagnostically Relevant Regions Retrieval. in International Conference on Medical Image Computing and Computer-Assisted Intervention (2019).

13. Mehta, N., Alomari, R. S. & Chaudhary, V. Content based sub-image retrieval system for high resolution pathology images using salient interest points. Conf. Proc. IEEE Eng. Med. Biol. Soc. 2009, 3719–3722 (2009).

14. Hegde, N. et al. Similar image search for histopathology: SMILY. NPJ Digit Med 2, 56 (2019).

15. Hemati, S. et al. CNN and Deep Sets for End-to-End Whole Slide Image Representation Learning. in Proceedings of the Fourth Conference on Medical Imaging with Deep Learning (eds. Heinrich, M. et al.) vol. 143 301–311 (PMLR, 07--09 Jul 2021).

16. Schaumberg, A. J. et al. Interpretable multimodal deep learning for real-time pan-tissue pan-disease pathology search on social media. Mod. Pathol. 33, 2169–2185 (2020).

17. Wang, J. Z. Pathfinder: multiresolution region-based searching of pathology images using IRM. Proc. AMIA Symp. 883–887 (2000).

18. Schaer, R., Otálora, S., Jimenez-Del-Toro, O., Atzori, M. & Müller, H. Deep Learning-Based Retrieval System for Gigapixel Histopathology Cases and the Open Access Literature. J. Pathol. Inform. 10, 19 (2019).

19. Kalra, S. et al. Yottixel - An Image Search Engine for Large Archives of Histopathology Whole Slide Images. Med. Image Anal. 65, 101757 (2020).

20. Riasatian, A. et al. Fine-Tuning and training of densenet for histopathology image representation using TCGA diagnostic slides. Med. Image Anal. 70, 102032 (2021).

21. Chen, C. et al. Fast and scalable search of whole-slide images via self-supervised deep learning. Nat Biomed Eng 6, 1420–1434 (2022).

22. Mormont, R., Geurts, P. & Maree, R. Multi-Task Pre-Training of Deep Neural Networks for Digital Pathology. IEEE J Biomed Health Inform 25, 412–421 (2021).

23. Wang, X. et al. RetCCL: Clustering-guided contrastive learning for whole-slide image retrieval. Med. Image Anal. 83, 102645 (2023).

24. Li, S. et al. High-Order Correlation-Guided Slide-Level Histology Retrieval With Self-Supervised Hashing. IEEE Trans. Pattern Anal. Mach. Intell. 45, 11008–11023 (2023).

25. Sayers, E. W. et al. Database resources of the national center for biotechnology information. Nucleic Acids Res. 50, (2022).

26. Komura, D., et al. Luigi: Large-scale histopathological image retrieval system using deep texture representations. *bioRxiv* (2018) doi:10.1101/345785.

27. Huang, Z., Bianchi, F., Yuksekgonul, M., Montine, T. J. & Zou, J. A visual-language foundation model for pathology image analysis using medical Twitter. Nat. Med. 29, 2307–2316 (2023).

28. Friedman, J. H., Bentley, J. L. & Finkel, R. A. An Algorithm for Finding Best Matches in Logarithmic Expected Time. ACM Trans. Math. Softw. 3, 209–226 (1977).

29. Bentley, J. L. Multidimensional binary search trees used for associative searching. Commun. ACM 18, 509–517 (1975).

30. Collins, I. & Workman, P. New approaches to molecular cancer therapeutics. Nat. Chem. Biol. 2, (2006).

31. Lu, M. Y. et al. Data-efficient and weakly supervised computational pathology on whole-slide images. Nature Biomedical Engineering 5, 555–570 (2021).

32. Howard, A. G. et al. Searching for MobileNetV3. ICCV 1314–1324 (2019).

33. Radford, A., et al. Learning Transferable Visual Models From Natural Language Supervision. *arXiv [cs.CV]* (2021).

34. Huang, G., Liu, Z. & Weinberger, K. Q. Densely connected convolutional networks. Proc. IEEE Comput. Soc. Conf. Comput. Vis. Pattern Recognit. 2261–2269 (2016).

35. Xu, H. et al. A whole-slide foundation model for digital pathology from real-world data. Nature 630, 181–188 (2024).

36. Chen, R. J. et al. Towards a general-purpose foundation model for computational pathology. Nat. Med. 30, 850–862 (2024).

37. Lu, M. Y. et al. A visual-language foundation model for computational pathology. Nat. Med. 30, (2024).

38. Chen, R. J. et al. Scaling vision transformers to gigapixel images via hierarchical self-supervised learning. in 2022 IEEE/CVF Conference on Computer Vision and Pattern Recognition (CVPR) (IEEE, 2022). doi:10.1109/cvpr52688.2022.01567.

39. Kather, J. N. et al. Predicting survival from colorectal cancer histology slides using deep learning: A retrospective multicenter study. PLoS Med. 16, e1002730 (2019).

40. Gamper, J., Alemi Koohbanani, N., Benet, K., Khuram, A. & Rajpoot, N. PanNuke: An Open Pan-Cancer Histology Dataset for Nuclei Instance Segmentation and Classification. in Digital Pathology 11–19 (Springer International Publishing, 2019).

41. Amgad, M. et al. Structured crowdsourcing enables convolutional segmentation of histology images. Bioinformatics 35, 3461–3467 (2019).

42. Amgad, M. et al. NuCLS: A scalable crowdsourcing approach and dataset for nucleus classification and segmentation in breast cancer. Gigascience 11, (2022).

43. Coudray, N. et al. Classification and mutation prediction from non-small cell lung cancer histopathology images using deep learning. Nat. Med. 24, 1559–1567 (2018).

44. Chakravarty, D., et al. OncoKB: A Precision Oncology Knowledge Base. JCO Precis Oncol 2017, (2017).

45. Liberzon, A. et al. The Molecular Signatures Database (MSigDB) hallmark gene set collection. Cell Syst 1, 417–425 (2015).

46. Jiang, P. et al. Signatures of T cell dysfunction and exclusion predict cancer immunotherapy response. Nat. Med. 24, (2018).

47. Li, Y. et al. Proteogenomic data and resources for pan-cancer analysis. Cancer Cell 41, 1397–1406 (2023).

48. Campanella, G. et al. Clinical-grade computational pathology using weakly supervised deep learning on whole slide images. Nat Med 25, 1301–1309 (2019).

49. Jiang, P. et al. Big data in basic and translational cancer research. Nature reviews. Cancer 22, (2022).

50. Douze, M., et al. The Faiss library. *arXiv [cs.LG]* (2024).

51. Malkov, Y. A. & Yashunin, D. A. Efficient and robust approximate nearest neighbor search using hierarchical Navigable Small World graphs. IEEE Trans. Pattern Anal. Mach. Intell. 42, 824–836 (2020).

52. Moffat, A. & Zobel, J. Self-indexing inverted files for fast text retrieval. ACM Trans. Inf. Syst. Secur. 14, 349–379 (1996).

53. Jégou, H., Douze, M. & Schmid, C. Product Quantization for Nearest Neighbor Search. IEEE Trans. Pattern Anal. Mach. Intell. 33, 117–128 (2011).

54. Gong, Y., Lazebnik, S., Gordo, A. & Perronnin, F. Iterative quantization: a Procrustean approach to learning binary codes for large-scale image retrieval. IEEE Trans. Pattern Anal. Mach. Intell. 35, 2916–2929 (2013).

55. Indyk, P., Motwani, R., Raghavan, P. & Vempala, S. Locality-preserving hashing in multidimensional spaces. in Proceedings of the twenty-ninth annual ACM symposium on Theory of computing 618–625 (Association for Computing Machinery, New York, NY, USA, 1997).

56. Rao, A., Barkley, D., França, G. S. & Yanai, I. Exploring tissue architecture using spatial transcriptomics. Nature 596, (2021).

57. Sundstrom, A., Grabocka, E., Bar-Sagi, D. & Mishra, B. Histological Image Processing Features Induce a Quantitative Characterization of Chronic Tumor Hypoxia. PLoS One 11, e0153623 (2016).

58. Subramanian, A. et al. Gene set enrichment analysis: a knowledge-based approach for interpreting genome-wide expression profiles. Proc. Natl. Acad. Sci. U. S. A. 102, 15545–15550 (2005).

59. Semenza, G. L. Targeting HIF-1 for cancer therapy. Nat. Rev. Cancer 3, (2003).

60. Hanahan, D. & Weinberg, R. A. Hallmarks of cancer: the next generation. Cell 144, (2011).

61. Saltz, J. et al. Spatial Organization and Molecular Correlation of Tumor-Infiltrating Lymphocytes Using Deep Learning on Pathology Images. Cell Rep. 23, (2018).

62. Yamauchi, M., Barker, T. H., Gibbons, D. L. & Kurie, J. M. The fibrotic tumor stroma. J. Clin. Invest. 128, (2018).

63. Murphy, K. M. & Weaver, C. Janeway’s Immunobiology: Ninth International Student Edition. (W.W. Norton & Company, 2016).

64. Reis, E. S., Mastellos, D. C., Ricklin, D., Mantovani, A. & Lambris, J. D. Complement in cancer: untangling an intricate relationship. Nat. Rev. Immunol. 18, 5–18 (2018).

65. Roumenina, L. T. et al. Tumor Cells Hijack Macrophage-Produced Complement C1q to Promote Tumor Growth. Cancer Immunol Res 7, 1091–1105 (2019).

66. Riihilä, P. et al. Tumour-cell-derived complement components C1r and C1s promote growth of cutaneous squamous cell carcinoma. Br. J. Dermatol. 182, 658–670 (2020).

67. Hanada, K. I., Yu, Z., Chappell, G. R., Park, A. S. & Restifo, N. P. An effective mouse model for adoptive cancer immunotherapy targeting neoantigens. JCI insight 4, (2019).

68. Weinstein, J. N. et al. The Cancer Genome Atlas Pan-Cancer analysis project. Nat. Genet. 45, (2013).

69. Grossman Robert L., et al. Toward a Shared Vision for Cancer Genomic Data. N. Engl. J. Med. 375, 1109–1112.

70. Kingma, D. P. & Ba, J. Adam: A Method for Stochastic Optimization. *arXiv [cs.LG]* (2014).

71. Kaufman, L. & Rousseeuw, P. J. Finding Groups in Data: An Introduction to Cluster Analysis. (John Wiley & Sons, 2009).

72. Shang, H. H. et al. Histopathology Slide Indexing and Search — Are We There Yet? NEJM AI 1, AIcs2300019 (2024).

73. Hafemeister, C. & Satija, R. Normalization and variance stabilization of single-cell RNA-seq data using regularized negative binomial regression. Genome Biol. 20, 296 (2019).

74. Robichaud, N., Sonenberg, N., Ruggero, D. & Schneider, R. J. Translational Control in Cancer. Cold Spring Harb. Perspect. Biol. 11, (2019).

75. Biswas, S. et al. IgA transcytosis and antigen recognition govern ovarian cancer immunity. Nature 591, 464–470 (2021).

76. De Martino, D. & Bravo-Cordero, J. J. Collagens in Cancer: Structural Regulators and Guardians of Cancer Progression. Cancer Res. 83, (2023).

